# Chromatin Remodelling in Damaged Intestinal Crypts Orchestrates Redundant TGFβ and Hippo Signalling to Drive Regeneration

**DOI:** 10.1101/2024.08.30.610472

**Authors:** Mardi Fink, Kizito Njah, Shyam J. Patel, David P. Cook, Vanessa Man, Francesco Ruso, Arsheen Rajan, Masahiro Narimatsu, Andreea Obersterescu, Melanie J. Pye, Daniel Trcka, Kin Chan, Arshad Ayyaz, Jeffrey L. Wrana

## Abstract

Cell state dynamics underlying regeneration are under-characterized. Intestinal damage prompts reprogramming into revival stem cells (revSCs) that reconstitute Lgr5+ intestinal stem cells (ISCs). Single nuclei multiomics of chromatin accessibility and transcriptomes during regeneration from irradition showed revSCs display epigenetic profiles shared with ISCs and differentiated lineages. Furthermore, while revSC genes are accessible throughout homeostatic epithelia, damage-induced global alterations in crypt and revSC chromatin converge on TGFβ, as well as Hippo pathways. We show TGFβ directly induces functional revSCs and demonstrate individual revSCs form organoids with reconstituted Lgr5+ ISCs. Despite this, loss of TGFβ signalling yielded mild regenerative defects. In contrast, interference in both Hippo and TGFβ abolished revSCs, precluded generation of new ISCs and led to rapid intestinal collapse. Thus, the epithelium is poised to engage the revSC regenerative program that relies on crypt-localized, transient morphogen cues that function in a compensatory manner to support intestinal regeneration.

## Introduction

The intestinal epithelium is highly dynamic, and continuously self-renews under homeostatic conditions. This is driven by *Leucine-rich repeat-containing G Protein coupled receptor 5* (*Lgr5*) positive intestinal stem cells (ISC) that are located in the base of the crypt of the small intestine and colon^1^. Lgr5+ ISCs are proliferative, undergo self-renewal, and give rise to both transit amplifying cells and all differentiated intestinal epithelial lineages. Despite the rapid turnover of the homeostatic intestine, Lgr5+ ISCs are highly sensitive to damage^2,3^. In response to injury these cells are lost and the intestinal epithelium is rewired to mount a regenerative response that reconstitutes Lgr5+ ISCs and re-instates homeostasis^4^. The molecular and cellular events underlying intestinal regeneration are not well understood. However, in response to damage, both the partially and terminally differentiated offspring of Lgr5+ ISCs demonstrate considerable plasticity and are proposed to dedifferentiate to Lgr5+ ISCs^5^. Single-cell profiling further revealed that reprogramming is associated with acquisition of a fetal-like signature^3,4,6^ that is manifested in a unique Clu+ cell that emerges in the crypt to give rise to new Lgr5+ ISCs^3,7,8^ that in turn reconstitute the epithelium. Although rare in the homeostatic adult intestine^9,10^, lineage-tracing showed Clu+ cells are multipotent, prompting their naming as revival stem cell (revSCs). The initiation of fetal programs has been linked to the Hippo pathway^11^, which acts through the co-transcriptional activator, Yap, to promote intestinal regeneration^4,12–14^, but how the intestinal epithelial genome is organized and interacts with signaling networks to direct crypt reprogramming events and regeneration remains unclear.

We performed multiomics profiling of the homeostatic and regenerating mouse intestine to simultaneously survey transcriptional output and altered chromatin accessibility. Chromatin accessibility associated with Lgr5+ ISC signature genes indicates a significant barrier to direct reprogramming, whereas revSC signatures are generally open and thus poised across the entire epithelium. Genome-wide analysis further showed damage-dependent reorganization of chromatin accessibility in the regenerative crypt, and a revSC genome structure that incorporated profiles of ISC and differentiated chromatin states with enrichment for Hippo and TGFβ-Smad elements. Although TGFβ effectively promotes formation of functional organoid-forming revSCs, interference with either TGFβ or Hippo signalling yields mild regenerative defects. However, knocking out both pathways leads to the complete loss of revSCs and their regenerative ISC offspring, thus leading to profoundly impaired intestinal regeneration. Our findings uncover how dynamic reorganization of the crypt genome converges with transient signalling in the crypt to generate a redundant system supporting tissue regeneration.

## Results

### Multiomic profiling of the damaged intestinal epithelium

We previously mapped the rapid kinetics of cell state changes in response to injury, and showed revSCs are relatively rare in the homeostatic intestine but are transiently induced in the crypt in response to 12 Gy gamma irradiation (IR)^15^. These studies also showed that revSCs are rare in the homeostatic and 1 dpi intestine, but begin emerging at 2 dpi, and peak at 3 dpi with subsequent conversion to Lgr5+ ISCs^4^. Therefore, to explore mechanisms underlying intestinal regeneration we focused on simultaneously profiling the transcriptomic (RNA-Seq) and genome accessibility (ATAC-Seq) in single nuclei of small intestinal epithelial cells when revSCs are emerging at 2 dpi, and for comparison, the undamaged (0 dpi) homeostatic intestine (**Figure 1a** and **Methods**). Integration of chromatin accessibility and transcriptomic information using Seurat’s weighted nearest neighbor (WNN) approach^16^ revealed 20 annotated WNN clusters (W0-W19) that included all the major cell types of the small intestinal epithelium (enterocytes-EC; Goblet-GC; Enteroendocrine-EE; Tuft-TC; and Paneth cells-PC), immune cells (CD8+ T cells), and a small number of fibroblasts (**Figure 1b**). Comparison of 0 and 2 dpi cell distributions showed that ECs displayed substantive damage-dependent changes due to shifts in crypt-villus zonation^4^ (see below), and GC, EE were similarly distributed (**Extended Data Fig. 1a, b**), while ISCs and revSCs were almost exclusively present at 0 or 2 dpi, respectively (**Extended Data Fig.1a, b**), similar to our prior observations^4^. The fetal stem cell marker *Ly6a*, as well as *F3* and *Atg9b* were robustly expressed in revSCs at 2 dpi, while *Clu* was expressed at lower levels, and *Anxa1* was sporadic (**Figure 1c**), which likely represents differing temporal kinetics of induction in emerging revSCs.

**Figure 1.**
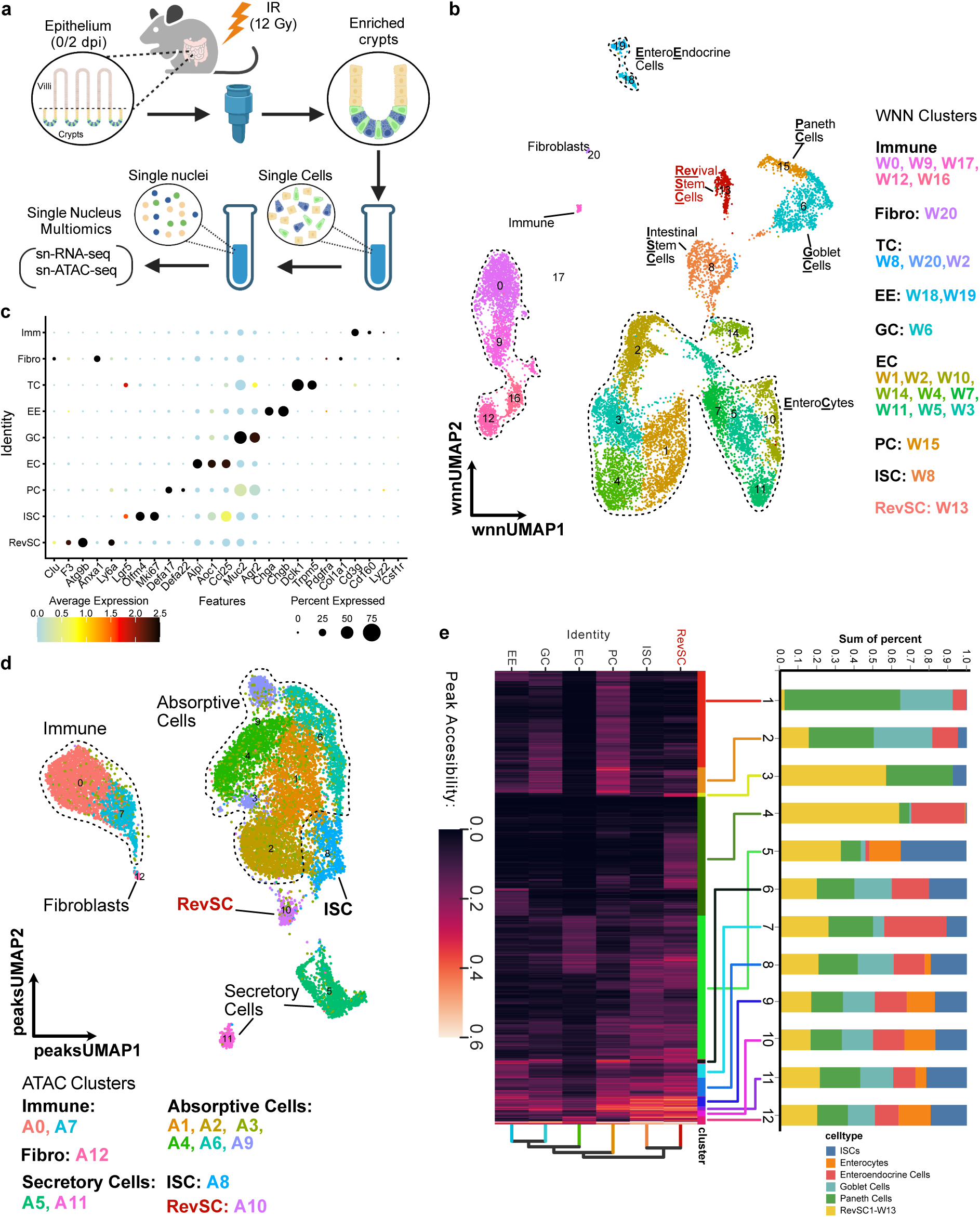
Multiomics survey of the regenerating fall intestinal epithelium. **(a)** Schematic of experimental design for ATAC and RNA-seq sample preparation from homeostatic and 2 dpi small intestines. **(b)** UMAP plot using unsupervised clustering of Weighted Nearest Neighbor (WNN) integration of single nucleus ATAC- and RNA-Seq data. Major cell types and their colour-coded clusters are indicated. **(c)** Dot plot of the average expression and percent of cells expressing the indicated gene in each of the major cell type clusters identified in **b**. **(d)** UMAP of the unsupervised clustering of single nuclei based on the ATAC-Seq data. Different clusters are colour-coded and cell identities assigned based on overlap of nuclei barcodes with WNN clusters. **(e)** Average peak accessibility across all cell types of the most significant differentially accessible peaks (Methods) associated with each of the indicated cell types was subjected to 2D hierarchical clustering and is plotted as a heatmap with scale was capped at 0.6 for visualization purposes. Peaks were clustered with cluster labels indicated by different colored bars **(right of heatmap)** and contributions of the indicated cell types to each cluster quantified **(bar graph)**.

Analysis of single nuclei transcriptomes showed similar patterns and cluster numbers as WNN (**Extended Data Fig. 1c, Table S1**), consistent with transcriptomes being an important driver of WNN clustering (**Extended Data Fig. 1d**). Interestingly, clustering ATAC profiles revealed only 13 clusters (A0-A12), and less diversity in the embeddings (**Figure 1d, Table S2**). To identify distinct cell types, we assessed how each WNN cluster contributed to the ATAC clusters (**Extended Data Fig. 1e**), which showed chromatin distinguished stem cells from differentiated cells, with the latter separated by their gross absorptive (A1, A2, A3, A6, A9) versus secretory commitment (A5, A8, A11) (**Extended Data Fig. 1e**). Furthermore, within the secretory lineage GC and PC were co-clustered in A5, while EE were found in A11, consistent with PCs and GCs arising from a common progenitor distinct from EEs^17–19^. Importantly, ISCs and revSCs were clustered in separate pre- and post-IR ATAC clusters (A8 and A10, respectively, **Extended Data Fig. 1e** and **f**), indicating that injury imparts significant, regeneration-associated shifts in chromatin accessibility in revSCs (**Extended Data Fig. 1f**). ATAC profiles associated with the major homeostatic intestinal lineages **(Table S3**) were then assessed for motifs revealing ISC peaks were enriched for WNT-associated transcription factors that mediate stemness, including TCF4 and KLF5^20^ (**Extended Data Fig. 1g, Table S4**). PCs also showed TCF4 motifs in agreement with their WNT-dependent maturation^21^. Secretory lineages were enriched for the fate determinant Atoh1, while EC ATAC peaks harbored Rxra and Nr2f6, which regulate EC gene expression^22,23^. We next explored if open chromatin strongly associated with specific lineages (904 peaks, **Table S5**) were accessible in other cell types by quantifying ATAC peaks across all cells (**Table S6**) and performing two-dimensional hierarchical clustering (**Figure 1e**). This showed most clusters contained peaks accessible in multiple cell types, supporting the notion of significant overlap in the genomic organization of otherwise phenotypically distinct intestinal cell types. However, some peak clusters unique to secretory cells were clearly evident (Cluster 1; **Figure 1e**), while Clusters 3 and 4 reflected peaks predominantly accessible in revSCs that were also accessible in EC and EE, respectively. RevSCs also displayed accessibility of heterogeneous ATAC clusters that were also open in both ISCs and a variety of differentiated cells (Clusters 5-12; **Figure 1e**). Since revSC can arise from the progeny of Lgr5+ ISCs and subsequently form fresh Lgr5+ ISCs^4^, these overlapping regions of genome accessibility may reflect manifestation of these dynamic cell state transitions. Thus, our dataset is well poised to interrogate the regulatory changes underlying both the homeostatic and early regenerating intestine.

### Mapping transcriptional and epigenetic dynamics of crypt stem cells and enterocyte zonation

Following injury, revSCs arise within the intestinal crypt, suggesting that cells along the crypt-villus (CV) axis may be differentially primed to become revSCs and/or exposed to spatially constrained signals following injury. To explore spatial dynamics of homeostatic and regenerative cell states, we used scVI^24^ to integrate transcriptomic profiles of revSCs, ISCs, and their EC progeny from 0 and 2 dpi cells (**Figure 2a**), omitting secretory cell lineages in order to focus on enterocyte differentiation dynamics in accordance with prior studies on crypt-villus patterns of zonation signature expression. We used Slingshot^25^ to infer a pseudotime trajectory (**Figure 2b**) that we subdivided into five discrete bins (Levels 1-5; **Extended Data Fig. 2a**) and found that in homeostatic conditions (0 dpi) Lgr5+ and Olfm4+ ISCs that reside in the base of the crypt were localized in Level 1 (**Figure 2c and d**). Analysis of EC zonation signatures (Clusters 1-5 from Moore *et al*.^26^) further showed Levels 1 through 5 captured crypt-villus spatial organization (**Figures 2c**; **Extended Data Fig. 2b**). Specifically, *Tfrc* (Moore Cluster 1) a crypt/early villus marker was expressed in Levels 1 and 2 and extended to Level 3 (**Figure 2c and Extended Data Fig. 2c**), where *Reg1* (Moore Cluster 2) expression, which peaks in early villus, was maximal (**Figures 2c and d**). This placed the CVJ at the boundary extending between Level 2 and 3. Accordingly, middle and late villus markers (e.g. *Slc2a2*, *Slc5a1* and *Slc15a1*, respectively) were expressed in corresponding gradients through Levels 3-5 (**Figures 2c, d**; **Extended Data Fig. 2c**) with Ada+ villus tip cells found exclusively in Level 5 (**Figures 2c, d**).

**Figure 2.**
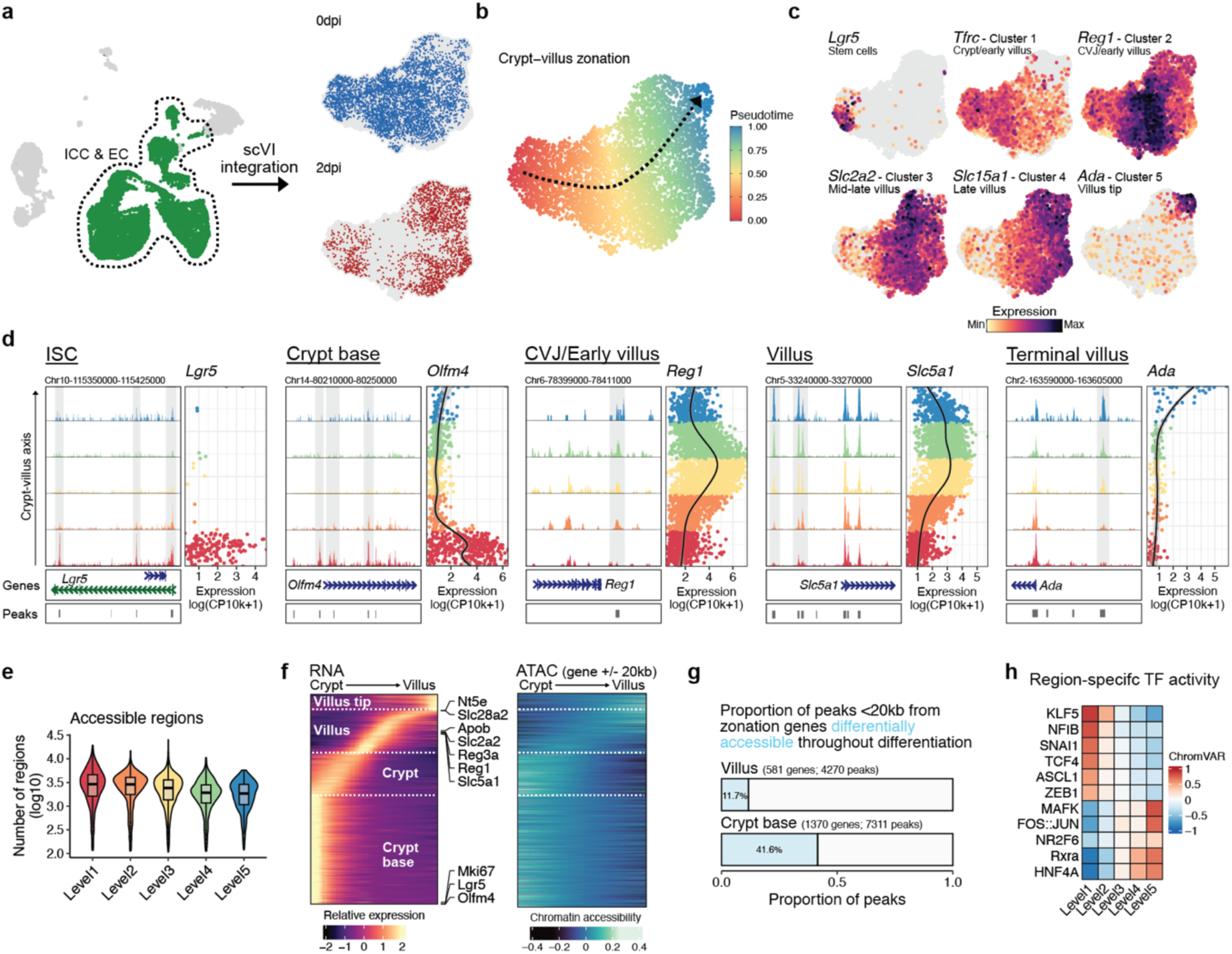
Transcriptional and epigenetic dynamics along the crypt-villus (CV) axis. **(a)** UMAP of snRNA-Seq data showing EC, ISC and revSCs clusters extracted for integration using scVI (left panel). Nuclei from either 0 dpi (blue), or 2 dpi (red) are plotted on the integrated UMAP (right panels) **(b)** Pseudotime values of an inferred continuous trajectory (dotted arrow) are colour coded and plotted on the integrated UMAP. **(c)** Integrated UMAP embeddings showing the relative expression of markers representative of the indicated zonation clusters identified by Moor et al. **(d)** Chromatin accessibility and expression of zonation markers along the inferred CV axis in 0 dpi cells. ATAC-Seq data of indicated loci are represented as the aggregated signal of cells within 5 discrete bins along the inferred trajectory (**see ED 2a**). Corresponding gene structure and ATAC-seq peaks are shown at the bottom of each panel. Grey bars highlight zonation-responsive peaks. Corresponding expression is shown and colour coded according to levels. For each gene, the height of ATAC traces were set to a common range and RNA expression is quantified as log-transformed UMI counts per 10k counts (log CP10k+1). **(e)** Total number of regions with accessible chromatin in cells along the CV axis in 0 dpi cells. **(f)** Modelled expression dynamics along the reconstructed CV axis in 0 dpi cells **(left)**, with genes ordered based on the location of their maximal expression (red heat scale), while corresponding chromatin accessibility is shown on the right (blue heat scale). **(g)** The proportion of peaks adjacent (<20kb) to zonation-associated genes that are dynamically accessible along the crypt-villus axis in 0 dpi cells. Zonation-associated genes were defined as those with differential expression (padj < 0.05, |log2FC| > 2) between crypt base cells (Level 1) and villus enterocytes (Levels 3/4). Peaks with a log2FC > 2 between regions were quantified. **(h)** ChromVAR activity scores for select transcription factors with motifs over-represented in differentially accessible loci associated with each level in 0 dpi cells.

To assess if regional expression was encoded in chromatin structure, we next assessed overall genome accessibility along the crypt-villus axis, which was relatively constant (**Figure 2e**). Furthermore, while transcript levels showed strong regional-specific expression, ATAC peaks associated with these genes were much less variable (**Fig. 2f**). Thus, zonation-associated enterocyte markers often harbored multiple regulatory elements that displayed preserved accessibility along the CV axis despite regionalized expression (unshaded peaks, **Figure 2d**, **Extended Data Fig. 2c**). In fact, of 581 genes upregulated during differentiation of enterocytes towards the villus tip (Level 3/4 vs. Level 1, log2FC >2, padj < 0.05), 88.3% of the 4270 adjacent peaks (<20kb from gene) were similarly accessible in crypt base cells (**Figure 2g**; examples in **Figure 2d** and **Extended Data Fig. 2c**). In contrast, the accessibility of 41.6% of peaks adjacent to genes selectively expressed in crypt base cells was lost upon differentiation (**Figure 2g**). In agreement, the stem cell marker genes *Lgr5* and *Olfm4* displayed prominent ATAC peaks in Level 1 that were largely absent in Levels 3 and 4 (**Figures 2d**). Similar to binding motifs mapped by cell type (**Extended Data Fig. 1e**), spatial mapping showed TCF4 was strongly enriched in Level 1 peaks, along with other fate determining factors (**Figure 2h, Table S7**), while nuclear receptor motifs (RXRA, HNF4A, NR2F6) found in enterocytes, showed increased accessibility at higher levels, and Level 5 cells showed evidence of accumulating stress (AP1 and MAFK; **Figure 2h, Table S7**). Collectively, these studies show that ECs do not undergo large-scale reorganization of chromatin accessibility, contrasting ISC genes that are rapidly condensed as offspring migrate up the crypt.

### RevSC emergence is associated with selective activity of Smad2/3 signaling in the crypt

Spatially restricted chromatin accessibility in ISCs may create a critical epigenetic barrier to de-differentiation that has important implications in understanding how ISCs are reconstituted following damage. We therefore investigated spatial dynamics and found a higher proportion of more differentiated, villus tip ECs at 2 dpi (**Figure 3a**), which is consistent with wide-scale loss of Lgr5+ ISCs (**Extended Data Fig. 1c**), as noted previously^4^. Differential expression analysis between 0 and 2 dpi cells further showed broad effects with maximal changes in Levels 2 and 3 (**Figure 3b**). Despite IR affecting multiple regions, expression of revSC markers, such as *Clu*, *Ly6a*, *S100a6*, *Cd44*, and others (**Table S8**) was restricted to the crypt regions (Level 1, p=1.1^e-7^; Level 2, p=9.6^e-7^; **Figures 3b,c**), and straddled Levels 1 and 2 (**Extended Data Fig. 3a**), consistent with their emergence in the crypt and CVJ^4^. To identify putative revSC regulatory mechanisms we evaluated global changes in spatial chromatin accessibility following IR by identifying differentially accessible peaks between 0 dpi and 2 dpi cells. Damage-dependent alterations were most abundant in the lower levels (**Figure 3d**), indicating cells in the crypt (Levels 1 and 2) display the most dynamic genomic responses to damage. However, when we examined revSC marker genes, most had accessible loci in 0 dpi cells, including components of the fetal signature (eg. *Anxa3, Clu, Ly6a*), despite minimal expression, and Lgr5 retained similar ATAC patterns in crypt cells (Levels 1 and 2), despite loss of Lgr5+ ISCs (**Figure 3e**; **Extended Data Fig. 3b**). Although transcriptional responsiveness of a marker at 2 dpi was often linked to gains in small numbers of elements near each locus (grey bars, **Figure 3e** and **Extended Data Fig. 3b**), only *Anxa1* and *Ly6d* showed exclusive accessibility in 2 dpi cells (**Extended Data Fig. 3b**). This suggests that regenerative fetal signatures are poised for activation and that regional signalling networks are the key mechanism driving induction of revSCs in the crypt. Accordingly, the Yap/Taz transcription factor target, Tead, which promotes induction of revSCs and fetal signatures, displayed a significant damage response in Level 1, but not in Levels 3-5 (**Extended Data Fig. 4a**).

**Figure 3.**
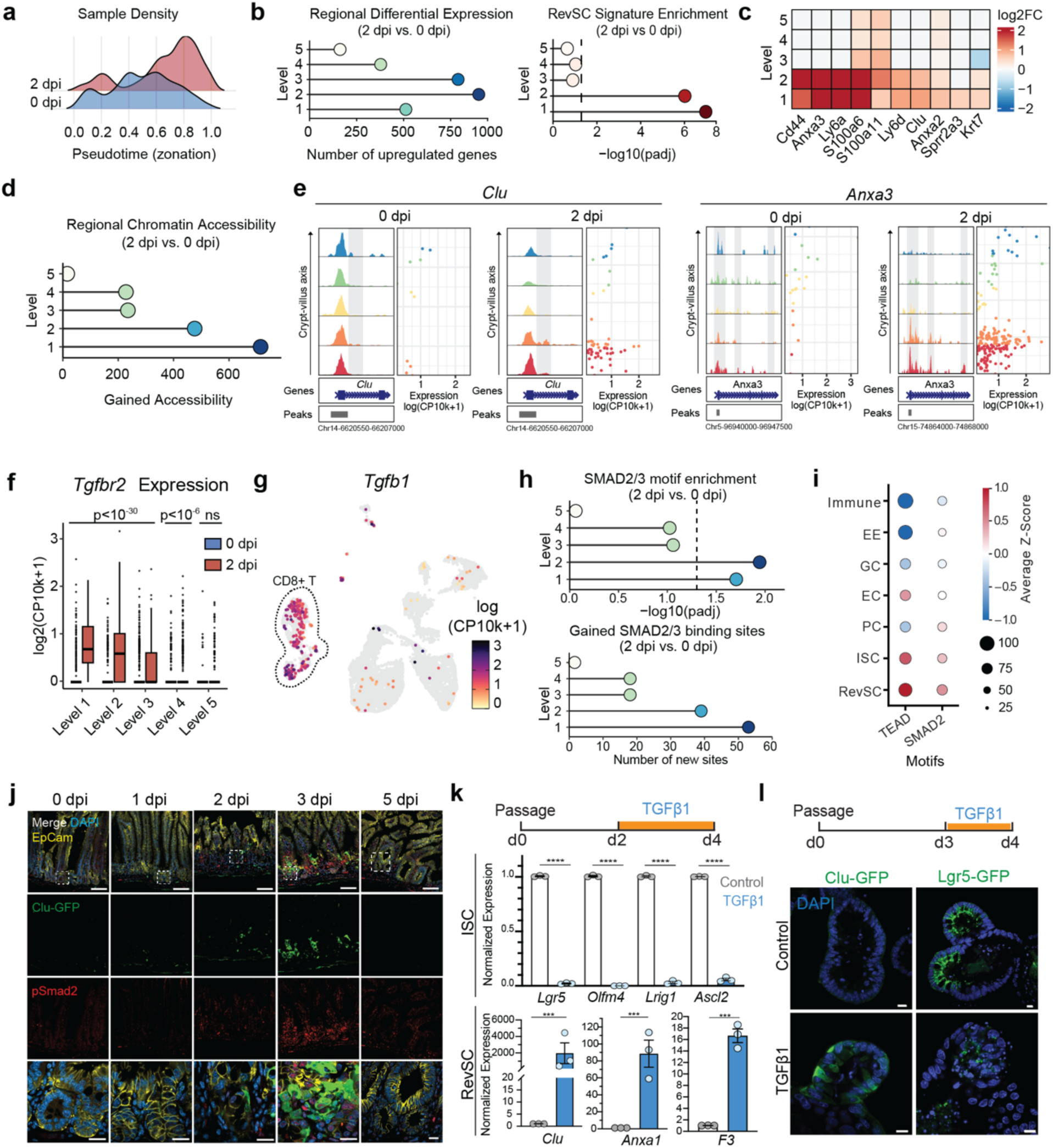
Spatial activation of TGFβ signalling in the regenerating crypt. **(a)** Relative distribution along the inferred trajectory of single nuclei from 0 dpi (blue) and 2 dpi (red) samples. **(b)** Number of genes differentially expressed (padj < 0.05, log2FC > 0.5) between 2 and 0 dpi cells within each of the five discrete regions is plotted according to the reconstructed CV axis **(left)**. Over-representation of the 50-gene revSC signature^4^ in the differentially expressed gene set of each region **(right)**. P-values were calculated using a Fisher’s exact test and corrected using the Benjamini-Hochberg method. Point colours reflect x-axis value. (c) 24312747v1Heatmap of log2 fold change in expression between 2 and 0 dpi nuclei of the indicated revSC genes along the CV axis. **(d)** Differential chromatin accessibility (padj < 0.05, log2FC > 0.5) between 2 and 0 dpi cells within the five discrete regions along the reconstructed CV axis. Point colours reflect x-axis value. **(e)** Chromatin accessibility traces and expression of revSC markers in 0 and 2 dpi cells along the CV axis. For each gene, the height of ATAC traces were set to a common range and RNA expression is quantified as log-transformed UMI counts per 10k counts (log CP10k+1). **(f)** *Tgfbr2* expression in 0 and 2 dpi cells along the reconstructed CV axis. Expression is quantified as log-transformed UMI counts per 10k counts (log CP10k+1) and plotted as box and whiskers (left panel; box ranges 25-75^th^ percentiles, bar represents median, and whiskers span 1.5x the interquartile range). **(g)** *Tgfb1* expression (log CP10k+1 counts; colour scale) in single nuclei is plotted (right panel) on the WNN UMAP. CD8+ T cells and immune cells are highlighted. **(h)** Smad2/3 binding motif (JASPAR: MA1964.1) over-representation (top) and motif counts (bottom) in differentially accessible peaks along the CV axis. **(i)** Dot plot indicating average ChromVAR Z-scores of motif enrichment within ATAC peaks accessible in each of the indicated cell types for TEAD and SMAD2. Size of dot indicates percentage of nuclei expressing their cluster’s Z-score average or greater/lesser depending on the sign of the average. **(j)** Immunofluorescence staining of pSmad2 (red), Clu-GFP (green) and EpCam (yellow) in wild-type mice following intestinal injury at the indicated dpi. *n=3* mice per condition. Scale bar, 100 μm. Boxes denote enlarged regions, scale bar 20 μm. **(k)** Experimental design **(top)** to assess ISC and RevSC signature genes expression **(bottom)** in control or TGFβ1-treated (200 pM; 48 h) WT organoids. Results from RT-qPCR were normalized against *Ywhaz* expression and are plotted as the mean (±SEM) of 3 independent experiments performed in triplicate. Unpaired, two-tailed t-test; ***p<0.0001. **(l)** Representative IF images of Clu-GFP **(left)** or Lgr5-GFP organoids **(right)** treated as indicated **(top, schematic)**, with TGFβ1 (200 pM; 24 h) and imaged for DAPI (blue) and GFP (green). Scale bar, 10µm.

To search for additional extrinsic cues promoting localized revSC production, we examined damage-induced genes along the crypt-villus axis (**Table S9**). This showed the TGFβ type II receptor, *Tgfbr2*, was one of the top damage-induced genes, particularly in Levels 1 and 2 (**Figure 3f**), with *Tgfbr2*-associated ATAC peaks concomitantly induced (grey bars, **Extended Data Fig. 4b**). On the other hand, expression of the TGFβ ligand, *Tgfb1*, was observed in immune cells (**Figure 3g**) that can migrate into damaged crypts^4^. We confirmed damaged-induced expression of *Tgfbr2* in crypt epithelia (**Extended Data Fig. 4c**), and expression of *Tgfb1* in Cd3e+ immune cells, Adgre1+ macrophages, and PDGFRa+ fibroblasts (**Extended Data Fig. 4d**). These results complement and confirm the recent identification of mesenchymal and immune sources of TGFβ1 in the regenerating intestine^27^. We also noted that expression of Smad2 and Smad3, which are the direct substrates of the TGFβ receptor, were elevated in Levels 1 and 2 (**Extended Data Fig. 4e**). Indeed, examination of transcription factor motif enrichment among injury-associated peaks (**Table S10**) showed that despite the diversity and low complexity of Smad motifs, we observed selective over-representation and gained sites in regenerating Level 1 and 2 crypt cells (p=0.02 and 0.01, respectively; **Figure 3h**). Furthermore, analysis by cell type showed Smad motifs were enriched in revSC-, ISC- and PC-associated ATAC peaks (**Figure 3i, Table S4**). Hippo signalling contributes to regeneration, and similar to Smads, sites harbouring the Yap transcription factor partner Teads were also enriched in these populations (**Figure 3i**). We validated these observations by staining for Smad2/3 protein, which was low at homeostasis (0 dpi), but by 2 and 3 dpi, was prominent in the crypt, and subsequently declined to baseline by 7 dpi (**Extended Data Fig. 4f**). Phosphorylation of Smad2 and Smad3 by the TGFβ receptor induces their nuclear accumulation. Therefore, we directly assessed TGFβ signalling activity by analyzing receptor-dependent phosphorylated Smad2 (pSmad2), which showed initiation of activation at 2 dpi with strong signals detected in the crypt by 3 dpi, when 188/243 (77%) of Clu+ cells displayed pSmad2 staining (**Figure 3j**). By 5 dpi pSmad2 was undetectable (**Figure 3j**). Finally, we analyzed inhibitory Smad7, which is a direct TGFβ-Smad target that functions in a negative feedback loop^28,29^. Smad7 protein and mRNA were low in homeostatic crypts, but by 2 dpi Smad7 protein was sporadic and by 3 dpi both protein and mRNA were broadly present in the crypt, returning to baseline by 5-7 dpi (**Extended Data Fig. 4g, h**). Collectively, these data demonstrate that in addition to Yap-Tead activation^30^, TGFβ signalling is transiently mobilized in the damaged crypt, corresponding with revSC induction, and both pathways are associated with the revSC genome.

To directly test if TGFβ induces revSCs, we next treated mouse intestinal organoids with TGFβ1, which potently induced the revSC markers *Clu*, *Anxa1*, and *F3*, and concomitantly suppressed ISC markers (*Lgr5, Olfm4, Ascl2, Lrig1*; **Figure 3k**). Similar TGFβ-dependent induction of revSC markers was recently reported^27^. Quantification of revSC induction using Clu-GFP organoids^4^, showed TGFβ dose-dependently activated Smad2/3 nuclear accumulation (**Extended Data Fig. 5a**) and induced Clu^+^ revSCs within 24h (**Figure. 3l**) that comprised about 1% of the population, as quantified using flow cytometry (**Extended Data Fig. 5b, c**). Of note, in control organoids, Clu+ cells were low but detectable and suppressed by treatment with the selective TGFβ receptor antagonist SB431542^31^ (SB, **Extended Data Fig. 5c**). Parallel analysis of organoids harbouring a Lgr5+-GFP allele showed TGFβ1 also potently suppressed Lgr5+ISCs and crypts (**Figure 3l**; **Extended Data Fig. 5d**). Thus, TGFβ signaling directly stimulates a regenerative phenotype by both inducing revSCs and suppressing Lgr5+ ISCs.

### TGFβ signalling induces functional revSCs

Damage by IR^4,32^, DSS^4,11^, or helminth infection^3,7^ depletes Lgr5+ ISCs that are restored via fetal-like regenerative populations^3,7,11^. To test if revSCs induced *in vitro* are functional, we treated organoids with TGFβ1, which did not substantially inhibit organoid numbers, but suppressed crypts that were reconstituted after a chase in the absence of TGFβ1 (**Figure 4a and Extended Data 5e**). To test if recovered crypts arose from revSC progeny, we performed lineage-tracing using *CluCreERT2;Rosa26-lsl-tdTomato* (Clu^tdTom^) organoids on a *Lgr5-GFP-DTR* background (hereafter Lgr5-GFP)^33^. To initiate tracing, *Clu^tdTom^;Lgr5-GFP* organoids were stimulated with TGFβ1 and 4-hydroxytamoxifen (4-OHT), prior to passaging and a 4 day recovery (**Figure 4b**). Quantification of tdTom-labelled crypts showed that 4-OHT treatment, independent of TGFβ1, led to a median of 18% traced Lgr5-GFP+ crypts (**Figure 4c**), consistent with mechanical stress from passaging inducing damage responses^34^. However, TGFβ1 strongly induced tracing, with mosaic patterns observed following 1 pM of TGFβ1 (median 34% of Lgr5+ regions) and robust (100%) clonal tracing observed at 50-200 pM of TGFβ1 (**Figures 4b**), while Lgr5-GFP+ crypt numbers were comparable across treatments (**Figure 4c**). Quantification of labelled cells by flow cytometry (**Extended Data Fig. 6a-c**) similarly revealed a dose-dependent increase in tdTom+ cells that reached ~90% of all cells, including Lgr5+ cells, at 50 and 200 pM TGFβ1 (**Fig. 4d**). Accordingly, immunofluorescence staining of the differentiation markers Muc2 (GC), ChgA (EE) and Lyz1 (PC) showed co-expression of tdTomato (**Extended Data Fig. 7a**), confirming that TGFβ1-induced Clu+ cells give rise to Lgr5+ ISCs and their differentiated progeny. TGFβ is a cytostatic factor that potently inhibits epithelial proliferation^35^. RevSCs are quiescent with low proliferation^4^, and we found a TGFβ1 pulse (24 h) suppressed proliferation in organoid crypts concomitant with induction of revSCs (**Figure 4e, left**), whereas proliferative, lineage-traced Lgr5+ crypts were noted after chasing out TGFβ1 with control media (**Figure 4e, right**). Further, TGFβ1-treated Clu+ cells are transient, as Clu+ cells were not maintained following passaging and removal of TGFβ1 from cultures, but were re-induced by 24 h treatment with TGFβ1 (**Figure 4f**).

**Figure 4.**
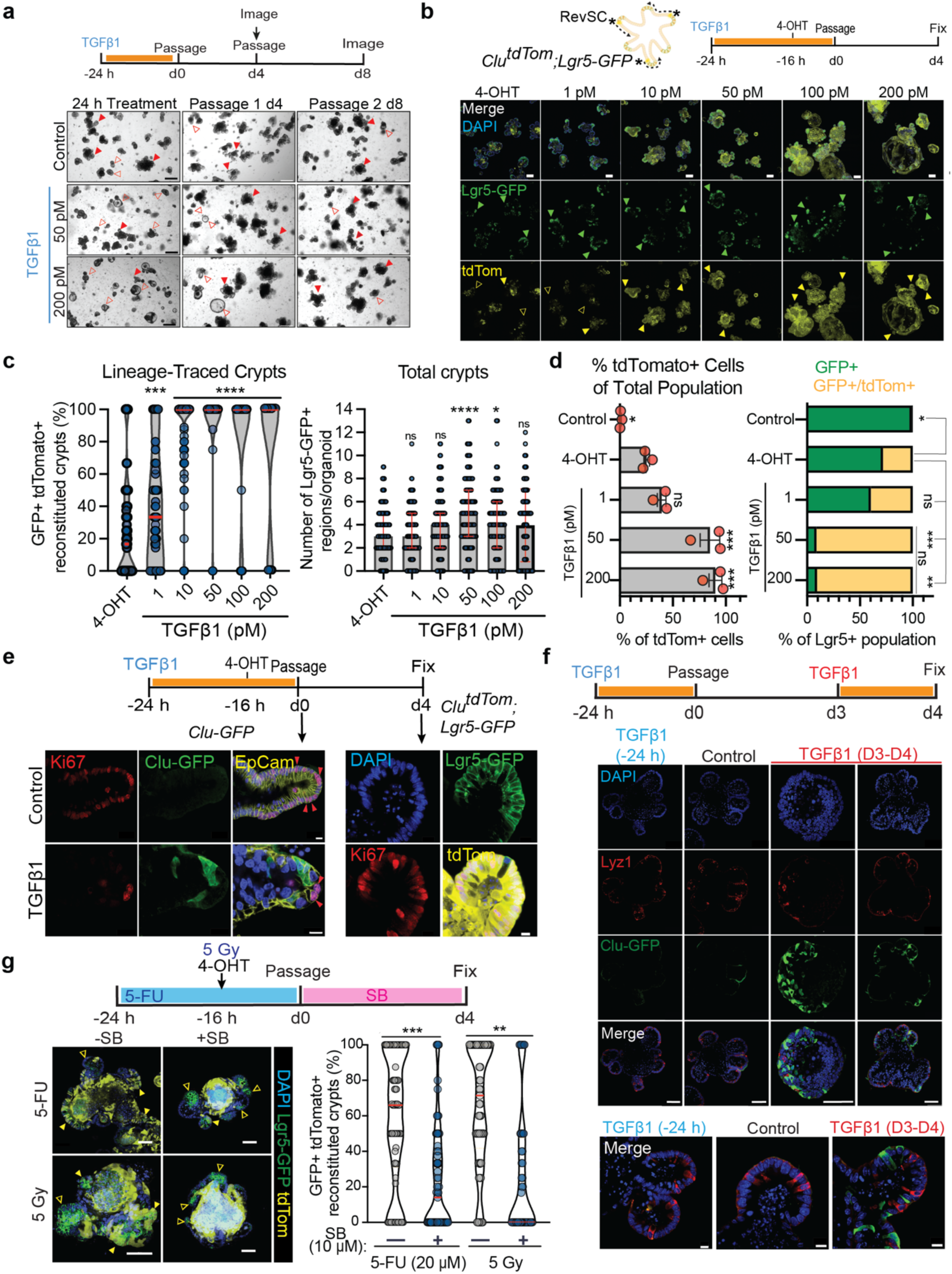
TGFβ signalling promotes the emergence of functional Clu+ revSCs. **a)** Schematic **(top)** and representative brightfield images of control or TGFβ1-treated (50 or 200 pM; 24 h) organoids. n=3 experiments, performed in triplicate. Solid red arrows indicate multiple crypts, open arrows indicate spheroids or single crypts. Scale bar, 500 µm. D, day; h, hour. **b)** *Clu-CreERT2;lsl-tdTom;Lgr5-GFP* organoids were treated according to the schematic with a pulse of TGFβ1 (24 h) and then 4-hydroxytamoxifen (4-OHT; 16 h) to induce recombination prior to passaging (top). Representative immunofluorescence images indicate Lgr5-GFP+ cells (green) and Clu-tdTomato+ progeny (yellow), nuclei stained with DAPI. Examples of GFP+ crypts (green arrows) that are either tdTom- (untraced; open yellow arrow) or tdTom+ (lineage-traced; closed yellow arrow) are highlighted. n= 2 independent experiments with each condition performed in triplicate. Scale bar,100 µm. **c)** Quantification for **(b)** of the percent of Lgr5+ regions that are lineage-traced (tdTom+) following TGFβ1-treatment represented as a violin plot, red lines indicate median. **(left panel)**. Total number of Lgr5-GFP+ regions formed after TGFβ1 treatment, bars represent median and interquartile range **(right panel)**. Significance calculated by unpaired, two-tailed t-test; *p<0.05, ***p<0.001, ****p<0.0001. **d)** Quantification from flow cytometry (See 4b schematic) of the percent of total tdTom+ cells. Bars represent the mean (± SEM) of n=3 individual experiments performed in triplicate (dots) **(left)**. The proportion of traced (tdTom+;GFP+; yellow) vs. untraced (GFP+;tdTom-); green Lgr5-GFP+ cells. Bars, represent the average proportion of cells over n=3 replicates **(right)**. Statistics: 2-tailed, unpaired t-test with 4-OHT-treated cells representing the comparison group. “Control” are organoids without 4-OHT treatment. ns= not significant, * p>0.05, ** p>0.01, *** p>0.001. **e)** Representative IF images of organoids untreated (Control) or treated with TGFβ1 (200 pM; 24 h; schematic) and stained for Ki67 (red), Clu-GFP (green) and EpCam (yellow), red arrows mark Ki67+ cells **(left)**. Organoids stained for Ki67 (red), Clu-tdTomato (yellow) and Lgr5-GFP (green) following a 4 day recovery after TGFβ1treatment **(right)**. Scale bar, 10 µm. D, day; h, hour. **f)** Representative IF images of Clu-GFP (green) organoids treated as indicated (schematic) and counter-stained with Lyz1 (red) to mark Paneth cells. Whole organoids are imaged **(top)**, scale bar, 50 µm. Crypt view **(bottom)**, scale bar, 10 µm. D, day; h, hour. **g)** *Clu-CreERT2;lsl-tdTom;Lgr5-GFP* were damaged with 5-FU (20 µM) or IR (5 Gy) and cultured in ENR media +/− SB431542 (SB; 10 µM) as indicated **(top)**. Representative IF images of Lgr5-GFP (green) and tdTomato (yellow). Solid yellow arrows indicate lineage-traced Lgr5+;tdTom+ crypts, open arrows depict Lgr5+;tdTom-crypts. Scale bar, 50 µM **(left)**. Lineage traced crypts in each condition were quantified and visualized in a violin plot **(right).** Red lines indicate median, n = 2-3 replicates. Unpaired, two-tailed t-test; *p<0.05, ***p<0.001.

**Figure 5.**
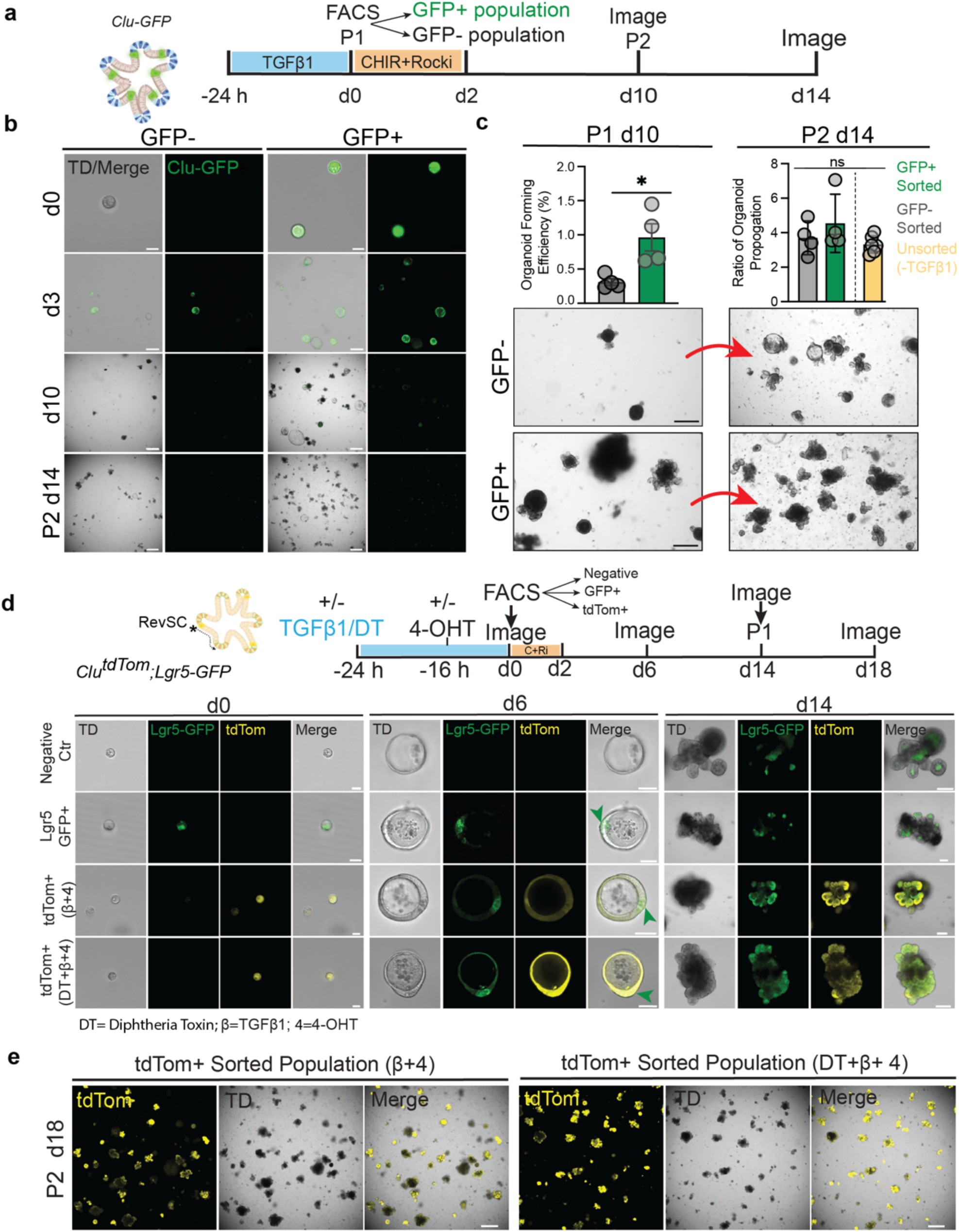
TGFβ-stimulated Clu+ cells are sufficient to generate organoid cultures. **a)** Schematic of organoid treatment prior to FACS and subsequent culture of GFP+/− cells from *Clu-GFP* organoids following treatment with TGFβ1 (24 h; 200 pM). P, passage number; d, day; h, hour; Rocki, Rock inhibitor (Y-27632); CHIR, CHIR 99021. **b)** Representative brightfield (TD) and fluorescent (GFP) images of sorted cells from *Clu-GFP* organoids at 0-, 3-, 10-, and 14-days in culture post sorting. Scale bar, 10 µm (with zoom) (d0); 100 µm (d3); 500 µm, (d10 and d14). **c)** Organoid forming efficiency of GFP+/− sorted cells. The indicated cells were plated at 10,000; 5,000; 1,000; or 500 cells/well and organoids formed after 14 days quantified and plotted as the mean ± SD of n=4 replicates **(left)**. Ratio of organoids propagated from P1 to P2 was quantified as organoids/ well for the indicated P1 cultures. Yellow bar denotes standard propagation efficiency in unsorted control cultures. Statistical significance via. Representative bright-field images of organoids derived from GFP- or + cells are shown **(bottom panel)**. Scale bar 295 µm. *p<0.05, unpaired, two-tailed t-test in **a** or one-way ANOVA in **b.** ns = not significant. **d)** Schematic of experimental design to assess organoid formation from individualized *Clu-CreERT2;lsl-tdTom;Lgr5-GFP-DTR*. Organoids were treated with or without TGFβ1 (β; 24 h; 200pM), Diphtheria toxin (DT; 24 h; 20 ng/mL) and/or +/− 4-OHT (16 h) and sorted as fluorescent negative, GFP+;tdTom- or GFP-;tdTom+ fractions **(top)**. Representative images are shown from brightfield (TD) and fluorescent channels (Lgr5-GFP; green, tdTom; yellow) of sorted cells after 0-, 6-, or 14-days in culture, as indicated. Green arrows denote emerging Lgr5+ cells (middle panels). Scale bar, 10 µm (with zoom) (d0); 50 µm (d6); 100 µm (d14). P, passage number; d, day. **e)** Representative images from TD and fluorescent channels of organoids from the tdTomato+ sorted populations obtained at 18 days post-sorting and imaged 4 days after passaging at day 14. Cultures derived from organoids treated with TGFβ1 and 4-OHT only (**left panel**) or TGFβ1, DT and 4-OHT (**right panel**). Scale bar, 500 µm. DT, Diphtheria Toxin; β, TGFβ1; 4, 4-OHT.

To test the importance of TGFβ signalling during *in vitro* regenerative signalling, organoids were treated with the DNA damaging agent, 5-Flourouracil (5-FU). This resulted in a loss of Lgr5-GFP+ crypts (**Figure 4g**, **Extended Data Fig. 7b**), with a return to normal morphology 8 days after recovery (**Extended Data Fig. 7c**). We also observed reduced expression of ISC markers (*Lgr5*, *Lrig1*, *Olfm4* and *Ascl2)* (**Extended Data Fig. 7d**), and induction of revSC markers (*Clu*, *F3* and *Anxa1*)(**Extended Data Fig. 7e**), To determine if revSCs reconstituted Lgr5+ ISCs during the recovery phase, we damaged Clu lineage-tracing organoids with either 5-FU, or 5 Gy IR, and allowed recovery, after which 66% and 71% of reconstituted crypts arose from revSC progeny (**Figure 4g**), with analysis of 5-FU-treated organoids confirming reconstitution of proliferative Lgr5+ISCs and their differentiated offspring (**Extended Data Fig. 5f**). Importantly, treatment with the TGFβ type I receptor inhibitor, SB431542, inhibited lineage-tracing in response to either damage cue, although regeneration was not completely blocked, as some traced and non-traced Lgr5+ crypts survived (**Figure 4g**). Finally, we examined if the regenerative pathways regulated by TGFβ in the mouse are conserved in humans using a colon organoid model (HCO, **Extended Data Fig. 8a**). Interestingly, HCOs are grown in the presence of the TGFβ receptor antagonist A83-01. Therefore, we examined if markers of mouse revSCs are regulated by TGFβ signalling by removing A83-01 and stimulating with TGFβ. This showed TGFβ potently suppressed crypts (**Extended Data Fig. 8a**), and analysis of mouse revSC markers (*CLU*, *ANXA1* and *F3*) further showed A83-01 removal induced expression of all three revSC marker genes that was strongly enhanced by ectopic TGFβ (**Extended Data Fig. 8b**). Furthermore, analysis of the direct TGFβ-Smad pathway target gene *SMAD7*, showed parallel inductions. Collectively, these results demonstrate TGFβ induces regenerative responses that may be conserved in human colon epithelia.

### Single TGFβ-induced revSCs form organoids

In the damaged intestine, we showed that revSCs reconstitute a new supply of Lgr5+ ISCs that drive regeneration, and here show that in organoids TGFβ induces functional revSCs that reconstitute Lgr5+ ISCs in response to damage. To test if isolated revSCs possess organoid-forming activity we therefore stimulated Clu-GFP+ organoids with TGFβ, isolated single Clu-GFP+ or Clu-GFP-cells by FACS and embedded them in Matrigel in the presence of CHIR and ROCK inhibitor for two days prior to transfer to standard EGF, Noggin and R-spondin containing media (**Figure 5a and Extended Data Fig. 9ai**), as previously described for propagating organoids from single Lgr5+ ISCs^36^. Immunofluorescence imaging of sorted cells immediately after plating (d0) confirmed isolation of GFP-versus GFP+ cells (**Figure 5b and Extended Data Fig. 9aii**). At 3 days post-sorting, while GFP+ cells retained Clu-GFP expression, some GFP-cells began expressing Clu-GFP+ (**Figure 5b**), in agreement with prior studies showing that passaging leads to induction of revSCs. By 10 days post-passaging, organoids with crypts were readily apparent in GFP+ sorted cultures and were abundant by day 14, with no detectable GFP (**Figure 5b**). Quantification showed that approximately 1% of Clu-GFP+ cells formed organoids (**Figure 5c**). In contrast, GFP-cultures formed fewer organoids (0.25% efficiency) and contained limited crypts until 14 days post-sorting, suggesting delayed reconstitution compared to GFP+ cultures (**Figure 5b, c**). Passaging of organoids from either condition, further showed efficient propagation similar to typical, unsorted organoids (**Figure 5c**).

To examine if sorted organoids arose from Clu+ cells, we next used the *CluCreERT2;lsl-tdTomato;Lgr5-DTR-GFP* lineage tracing model (**Figure 5d**) and sorted either Lgr5-GFP+ cells from untreated cultures, or tdTom+ cells from TGFβ1 and 4-OHT treated cultures (**Figure 9b**). Immediately after sorting we confirmed isolation of Lgr5-GFP+;tdTom-versus Lgr5-GFP-;tdTom+ single cells from untreated, or TGFb-treated cultures, respectively (**Figure 5d, second versus third row**). By 6 days in culture, Lgr5-GFP+ spheroids were apparent in both conditions, and by d14 both untreated and TGFβ-treated single cells generated organoids containing Lgr5-GFP+ crypts (**Figure 5d**). Of note, Lgr5+ cells that formed from isolated tdTom+ cells were lineage-traced confirming their origin from Clu+ revSCs. We note in these studies that Lgr5-GFP-cells also formed Lgr5+ GFP organoids (**Figure 5d**, **top row**), which likely reflects regenerative responses induced upon passaging of organoids^34^. To exclude the possibility that Lgr5+ ISCs might also express Clu, and thus provide the organoid-forming activity in Clu-sorted cells, we also ablated Lgr5+ cells prior to sorting using diphtheria toxin (DT) (**Figure 5d, bottom row**). However, despite cell ablation, Lgr5-GFP+;tdTom+ spheroid structures that were indistinguishable from non-DT treated cultures were apparent a d6, and by d14 DT-treated single cells had formed crypt-containing organoids, similar to non-DT treated conditions (**Figure 5d**). Moreover, organoids derived from revSCs either in the absence or presence of DT were readily passaged and gave rise to tdTom+ organoids (**Figure 5e**). These studies thus demonstrate that TGFβ-induced single Clu+ revSCs form organoids in the absence of Lgr5+ ISCs via reconstitution of a fresh supply of crypt ISCs.

### Interfering with TGFβ signalling yields moderate regenerative defects

To determine if TGFβ contributes to *in vivo* regenerative signaling we assessed the impact of *Tgfβr2* loss on regenerative signalling by crossing *Vil-CreERT2;Tgfbr2^f//fl^* mice (*Tgfbr2^−/−^*) to our previously generated Clu-GFP line, and confirmed efficient TAM-dependent knockout (**Extended Data Fig. 10a and b**). While homeostatic intestines appeared normal, we observed a reduction in Clu-GFP+ revSCs at 3dpi (35 versus 22 Clu+ cells/1000um, p<0.001; **Extended Data Fig. 10c**). To confirm loss of TGFβ signalling we assessed Smad2/3 induction, nuclear localization, pSmad2, and *Tgfbr2* expression levels. Increased Smad2/3 protein observed at 2 and 3 dpi was reduced in the absence of *Tgfbr2* in both the proximal and distal region of the small intestine (**Extended Data Fig. 10d**), consistent with early studies showing TGFβ induces Smad2 expression^37^. pSmad2 staining, assessed at 3 dpi when TGFβ signalling peaks, was also reduced upon loss of *Tgfbr2* (**Extended Data Fig. 10e**), and we confirmed by smFISH that the upregulation of *Tgfbr2* observed in damaged crypts was absent in the epithelium of the knockouts (**Extended Data Fig. 10f**). We also noted that TGFβ pathway activation was similar across the proximal-distal axis of the small intestine (**Extended Data Fig. 11a**), as was Clu+ cell production and its reduction upon *Tgfbr2* loss (**Extended Data Fig. 11b**). Finally, tracking the emergence of regenerative crypts showed that *Tgfbr2^−/−^*intestines had reduced Olfm4+ crypts at 5 dpi, but had substantially recovered by 7 dpi (**Extended Data Fig. 10g**). Thus, while TGFβ signalling is mobilized in the crypt, and TGFβ directly induces revSCs, interference with the pathway leads to moderate regenerative defects.

### The TGFβ and Hippo pathways cooperatively drive intestinal regeneration

The delay in crypt regeneration and modest phenotype in *Tgfbr2* mutant intestines is reminiscent with previous studies from us and others on the role of Hippo signalling in intestinal regeneration^15^, which showed in Yap and Yap/Taz knockouts that while intestinal regeneration in response to acute injury is impaired, intestines eventually recover^7,12,14,38^. We therefore examined the impact of Yap/Taz loss on revSC production. We confirmed Yap/Taz activation in both the proximal and distal small intestine **(Extended Data Fig. 12a)**, and found that Clu+ cells in Yap/Taz-deficient intestinal epithelia were reduced to ~17% of WT at 3 dpi (~50 versus 300 Clu+ cells/mm, **Extended Data Fig. 12b**) with concomitant delayed recovery of Olfm4+ crypts **(Extended Data Fig. 12c**), similar to our observations in *Tgfbr2*-deficient intestines. Similarly, infection with the symbiotic parasite *Heligmosomoides polygyrus bakeri* (Hbp) showed revSC production was only partially dependent on Yap^7^. The partial requirement for either *Tgfbr2* or *Yap* during intestinal regeneration suggested that transient activation of these pathways may cooperatively regulate the regenerative response. Indeed, our analysis of chromatin accessibility and DNA binding motifs in the crypt showed both the Yap target Tead, and Smad2/3 sites were concurrently induced by damage at 2 dpi and were enriched in revSCs (**Figures 3h, i**). Since TGFβ-Smad and Hippo-Yap-Tead pathways interact in various contexts^39^, we assessed Smad2/3- and TEAD-containing peaks and found that 28% of revSC-associated peaks contained these motifs (734 of 2612 peaks; p< 10^−70^; **Figure 6a**, **Tables S11**), and of these, most were either Tead or Smad, with only 48 containing both motifs. This suggests that Smads and Yap predominantly associate independently with the revSC genome. To determine Smad2/3 and Tead activity across the epithelium, we next extracted Smad2/3-only, Tead-only, and Smad-Tead composite peaks from the global ATAC peak set (**Table S12**) and calculated the mean accessibility per nuclei in each cell type (**Figures 6b**). This showed crypt-resident populations (revSC, ISCs, and Paneth cells) displayed elevated accessibility for all three classes when compared to the rest of the epithelium, with revSCs displaying the highest-level occupancy of all three classes (average p<2×10^−7^). Furthermore, their presence in ATAC profiles decreased in villus bottom and middle ECs. These results show that Yap/Taz and TGFβ-Smad signaling both converge on the revSC genome and may cooperate to redundantly drive regeneration.

**Figure 6.**
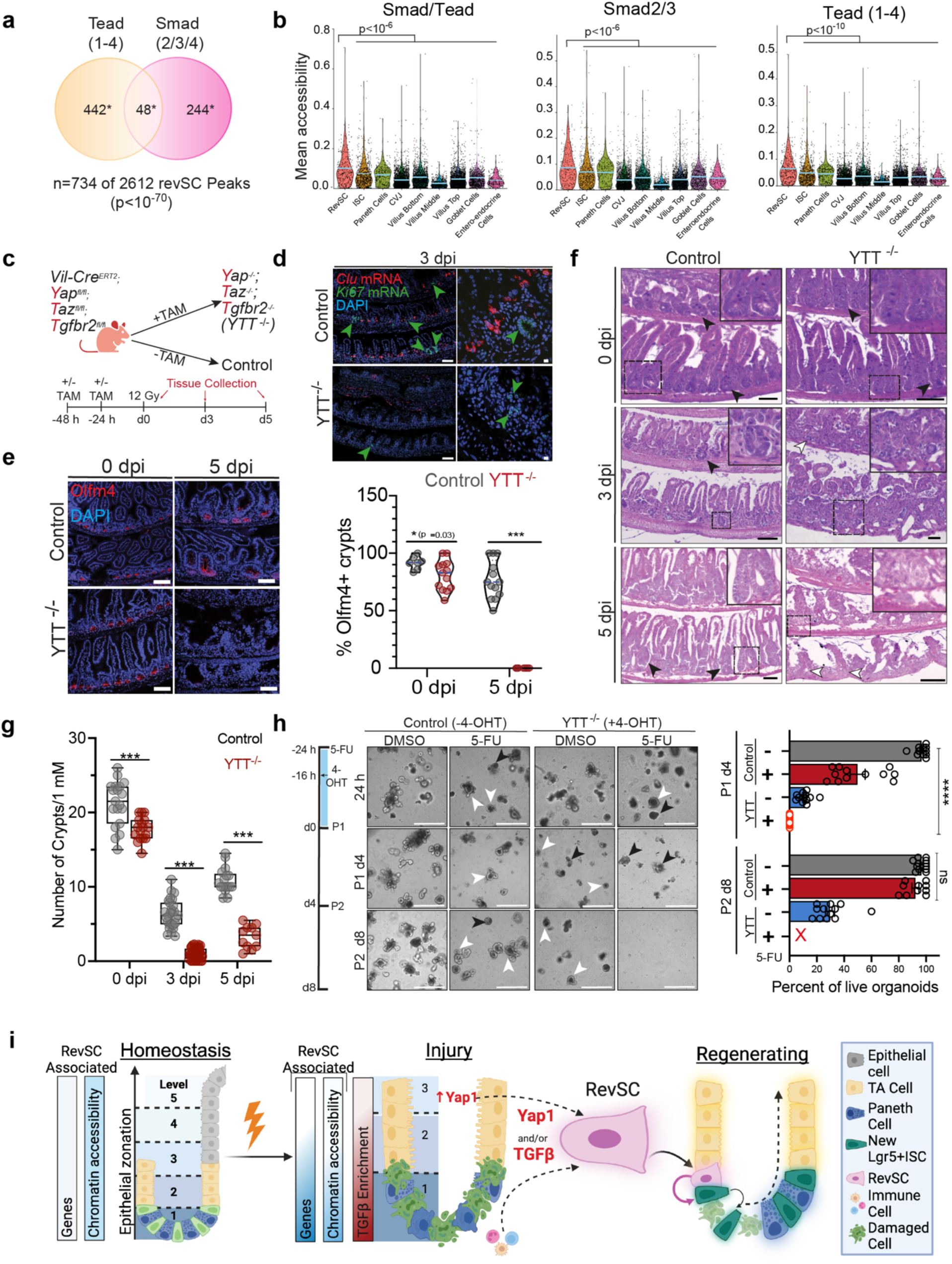
TGFβ and Hippo form a redundant system driving regeneration. **a)** Venn diagram measuring the overlap between Tead- and Smad-containing revSC peaks. p values for Smad alone, Tead alone and Smad-Tead overlap are <10^−16^, 10^−40^, and 10^−10^, respectively; Fischer’s exact t-test. **b)** Mean accessibility of Smad2/3-Tead peaks in the global peak set was quantified in single nuclei from the indicated cell types and visualized in a violin plot **(left)**. Mean accessibility of peaks from the total peak set that have a Smad2/3 site **(middle)** or a Tead 1-4 site **(right)** in each of the indicated cell types of the small intestinal epithelium. Statistical significance via unpaired, two-tailed t-test. **c)** Schematic of *Vil-CreERT2;Yap ^fl/fl^;Taz ^fl/fl^;Tgfbr2^fl/fl^* mouse model and experimental design **(6d-6g)**. Control (-TAM) and YTT^−/−^ (+TAM). **d)** RNAscope for *Clu* (red) and *Ki67* (green) expression at 3 dpi in control or YTT^−/−^ mice. Green arrows indicate Ki67+ regions. A rare example of two Ki67+ cells found in YTT^−/−^ intestines is shown. Scale bar, 100 µm (left); scale bar, 10µm (right). **e)** Representative IF images of Olfm4 (red) in control or YTT^−/−^ mice at homeostasis (0 dpi) or 5 dpi. Scale bar, 100 µm **(left)**. Quantification of percentage Olfm4+ crypts indicated in a violin plot, blue line indicates median **(right)**. Statistical significance calculated using unpaired, two-tailed t-test; *p<0.05 ***p<0.001. *n=2* mice per condition. **f)** Representative images of H&E staining of control or YTT^−/−^ knockout mice at 0, 3, and 5 dpi. Dashed boxes denote enlarged regions. Dark arrows depict visible crypts. Scale bar, 100µm. *n*=2 mice per genotype/condition. **g)** Quantification of the total number of crypts per 1mm of the small intestine. Statistical significance calculated by unpaired, two-tailed t-test; *p<0.05; 0. ****p<0.0001. Whiskers show maximum and minimum values, box extends from the 25^th^ to 75^th^ percentiles, with median represented by the center line. **h)** *Yap^fl/fl^;Taz^fl/fl^;Tgfbr2^fl/fl^*organoids either untreated (Control) or treated with 4-OHT (YTT^−/−^) and/ or 5-FU prior to passage as shown in the schematic. Representative phase contrast images were acquired at 24 h following treatment, and prior to passage 1, d4 and passage 2 d8. White and black arrows denote live and fragmented, dead organoids, respectively. *n*=3 experiments performed in triplicate. Scale bar, 500 μm **(right)**. Quantification of the percent of live organoids after passage 1 and 2. “X” indicates no detected organoids. Bars represent the mean (± SEM) **(left)**. Statistical significance via unpaired, two-tailed t-test; ****p<0.0001, ns=not significant. **i)** Model of changes in genome accessibility and transcriptional output related to revSC generation, highlighting the role of redundant TGFβ and Yap signalling in the conversion from revSCs to fresh Lgr5+ ISCs, which reconstitute the intestine following injury.

To test if these pathways compensate for each other, we generated *Vil-CreERT2;Yap^fl/fl^;Taz^fl/fl^;Tgfbr2^fl/fl^* (YTT) mice to direct inducible triple knock out (YTT^−/−^) intestinal epithelia following Tamoxifen treatment (**Figure 6c**). Similar to both *Yap;Taz*^12^- and *Tgfbr2*-deficient intestines, under homeostatic conditions mice appeared healthy 5 days after inducing YTT loss, and intestinal morphology was normal (see **Figure 6f**), despite a slight decrease in crypt density. However, in response to 12 Gy irradiation, TAM-treated YTT−/− mice rapidly declined, were moribund by 4 dpi, and required sacrifice by 5 dpi. Analysis of Clu+ revSCs by smFISH at 3 dpi showed typical induction in non-TAM treated controls, with regenerating proliferative crypts (*Mki67+*) evident (**Figure 6d**). In stark contrast, Clu+ revSCs were undetectable in the epithelium of TAM-treated YTT−/− intestines, with little to no proliferative crypts (**Figures 6d**). Analysis of Olfm4+ crypts in YTT−/− homeostatic intestines showed a slight reduction compared to normal homeostatic intestines and in response to damage Olfm4+ crypts were lost in both WT and YTT−/− crypts at 3 dpi, as expected (**Figure 6e**). In stark contrast, whereas at 5 dpi abundant, reconstituted Olfm4+ crypts were apparent in regenerating controls, YTT TKO intestines displayed no detectable Olfm4+ staining (**Figure 5e**). Consistent with the profound loss of regenerative signalling, gross morphology of the YTT TKO intestines showed small, degenerating crypts at 3 dpi, and by 5 dpi a dramatic loss of villi and villus architecture, with few if any detectable crypts (**Figure 6f and g**). Next we generated organoids from *Vil-CreERT2;Yap^fl/fl^/;Taz^fl/fl^;Tgfbr2^fl/fl^* intestines and tested response of YTT^−/−^ organoids to 5-FU-induced damage (**Fig. 6h**). In the absence or presence of 4-OHT, 5-FU induced loss of crypts during the acute phase of the damage response, with 4-OHT untreated control organoids subsequently recovering, similar to our studies above (**Extended Data Fig. 7c**). However, 5-FU-treated YTT−/− organoids failed to recover crypts, displayed a fragmented morphology, and yielded no viable organoids upon subsequent passaging (**Fig. 6h**). This dramatic collapse of organoids in response to damage contrasts the capacity of organoids to accommodate inhibition of either TGFβ (**Figure 4g**), or Yap/Taz signalling^14,30^. We also observed that passaging of 4-OHT-treated YTT−/−organoids led to declining organoid numbers, consistent with passaging inducing regenerative responses, as noted above (**Figure 6h**). Altogether, these findings show that revSCs are co-regulated by TGFβ and Hippo signalling that function in parallel pathways to promote induction of revSCs and intestinal regenerative responses (**Figure 6i**).

## Discussion

Intrinsic regenerative capacity is essential to facilitate the return to physiological homeostasis following acute injury. The intestinal epithelium is highly dynamic and in homeostasis is rapidly renewed via crypt-resident Lgr5+ ISCs, with distinct EC, secretory lineage and even ISC subpopulations patterned regionally along the crypt villus-axis^40^. This heterogeneity likely contributes to the highly adaptable responses of the gut to perturbations, which includes broad activation of some fetal signature genes across damage models^3,4^, as well as pathogen-specific shifts in cell proportions and variable activation of immune and stress response programs^40^. Our previous work identified revSCs, a regenerative population that expresses aspects of the fetal signature and is induced following various forms of injury that include IR, Lgr5+ ISCs ablation, DSS colitis, and parasites^4,7,41^. Here we provide deep single nuclei multiomics profiling of the homeostatic and irradiated intestinal epithelia that identifies important crypt-associated chromatin dynamics and identifies a critical role for parallel TGFβ and Hippo convergence on the revSC genome that provides a redundant system that robustly supports intestinal regeneration.

We found that while transcriptome dynamics were strongly associated with location, alterations in ATAC patterns were less variable, for example ATAC patterns in ECs undergoing crypt-villus state transitions were highly similar, and PC and GCs, which arise from a common progenitor^17,19^ clustered together and shared accessibility profiles. These findings indicate that spatially constrained cell states along the crypt-villus axis are not encoded in differential chromatin accessibility and that differentiated cell states are dependent on extrinsic cues that fine tune the transcriptomic output to drive sub-specialization. ISCs however are different, as ISC signature genes displayed unique ATAC peak profiles that were restricted to the crypt base, suggesting that chromatin structure represents a significant barrier to damage-induced direct reprogramming. In contrast, while a handful of revSC signature genes showed damage-dependent chromatin accessibility (e.g., *Anxa1*), most revSC signature genes were open across the crypt-villus axis, making them available for damage-dependent signalling. Despite this, damage-associated changes in chromatin were localized to the crypt, and our extensive analysis of TGFβ pathway activity also showed strong, transient activation in the crypt that corresponded to where and when revSCs emerge. This restriction of TGFβ signalling is likely manifested by localized sources of TGFβ that we and others show is produced by immune cells, macrophages and fibroblasts^30^.

TGFβ has previously been implicated in intestinal repair, with evidence suggesting that it might promote regeneration, but it is also linked to intestinal pathologies such as IBD, tumorigenesis and fibrosis^42,43,44,45,46^. Three recently posted studies similarly showed TGFβ induces fetal-like genes in intestinal models, with one identifying revSC and fetal characteristics in a CRC context, drawing implications between pathways that regulate regenerative and pathological programs ^27,47,48^. In agreement Yap-dependent induction of regenerative pathways is required for adenoma formation^30^. Moreover, the strong suppression of Lgr5+ ISCs further suggests TGFβ activity can control the balance between fetal-like revSCs and adult homeostatic ISCs.

Systematic screening of morphogen networks in organoids has shown complex pathway interactions mediate cell state dynamics in the epithelia^49^. Since Yap activity is also transiently induced in the damaged crypt similar to TGFβ^4,12^ and both promote revSC production, we posited the mild phenotype displayed by *Tgfbr2* intestinal knockout mice following injury reflected convergent regulation of revSCs (**Figure 6i**). Thus, while regeneration is possible in the absence of either TGFβ or Hippo signaling^12,14,38^, the catastrophic intestinal failure seen in the absence of both pathways demonstrates that independently perturbing either pathway makes the alternative pathway essential for regeneration. Accordingly, we found that Yap and Smads predominantly converge on distinct regions of chromatin in revSCs. This compensatory mechanism may explain why loss of Yap in parasitic infections results in only a partial loss of revSCs^40^. Of note, a recent publication linked intestinal epithelial fetal-reversion to TGFβ through a stepwise mechanism, in which TGFβ activates YAP/TEAD and SOX9 signaling, whereas our molecular and genetic analyses *in vivo* and in organoids indicate a redundant system^27^. Thus, further investigations are required to unravel this complex interaction and how these pathways may function either independently or as part of an orchestrated mechanism to regulate intestinal regeneration.

Extensive investigation of the homeostatic versus damaged intestine has uncovered key roles for de-differentiation in driving recovery, but the chromatin dynamics and signaling pathways governing reprogramming are only poorly understood. Analysis of chromatin accessibility suggests a barrier to rapid, direct reprogramming to ISCs, and we show that the revSC chromatin exists in an intermediary state that contains both differentiated and ISC ATAC profiles. Moreover, we now show that isolated TGFβ-induced revSCs can form organoids and reconstitute the Lgr5+ ISC pool, clearly demonstrating their regenerative capacity. These findings raise the question as to why de-differentiation would require an intermediary state to drive reconstitution of Lgr5+ ISCs. We recently showed that transient p53 activity is also critical for the emergence of revSCs during intestinal regeneration, adding a third pathway that controls revSCs^50^. Multiple pathways cooperating to induce revSCs might constitute a regenerative checkpoint in which signals must temporally and/or spatially converge to induce fit, fetal-like revSCs that then reconstitute healthy Lgr5+ ISCs thus protecting the intestinal genome from deleterious mutations. It will be of interest to determine if this putative regenerative checkpoint also functions in other tissues subject to frequent cycles of injury and repair.

## Acknowledgements

The authors thank the Network Biology Collaborative Centre (<u>nbcc.lunenfeld.ca</u>) for single cell profiling and sequencing service. The NBCC is a facility supported by Canada Foundation for Innovation, the Ontario Government, Genome Canada and Ontario Genomics (OGI-139). We thank L. Attisano (University of Toronto) for critical comments. This research is supported by the Canadian Cancer Society (grant #706715) and the Terry Fox Research Institute (Program Project Grant #1107) to JLW. Schematics throughout the manuscript were created with Biorender.com.

## Author Contributions

A.A., M.F., and J.L.W co-conceived the Study; A.A. and D.T designed and executed the Multiomics experiment. K.C. performed 10x genomics. S.J.P and D.P.C performed all Multiomics data processing and analysis, D.P.C. applied scVI data integration methods. M.J.P. maintained mouse colonies for various genotypes involved, along with D.T. and K.N. M.F performed experiments pertaining to *Tgfbr2^−/−^* mice, M.F and K.N. performed experiments pertaining to YTT mice; this included tamoxifen and irradiation treatments, tissue harvesting, preparation and characterizations by H&E, IHC, IF staining and RNAscope. V.M. and F.R. aided in additional tissue preparation and processing. M.F. performed *in vitro* organoid experiments. M.F., S.J.P, D.P.C., K.N, and J.L.W. wrote the manuscript.

## Methods

### RESOURCE AVAILABILITY

#### Lead contact

Further information and requests for resources and reagents should be directed to and will be fulfilled by the lead contact, Dr. Jeffrey Wrana

#### Materials availability

New experimental mouse models were developed for this study, as described below under “experimental model and subject details”.

#### Data and code availability

Multiomic data have been deposited at Geo and are publicly available as of the date of publication. All original code has been deposited at Zenodo and is publicly available as of the date of publication.

### EXPERIMENTAL MODEL AND SUBJECT DETAILS

#### Mice

Crl:CD1 (ICR) mice used as wild-type controls were provided by The Centre for Phenogenomics (Toronto). *Clu-CreERT2* was generated in-house as previously reported^4^. *Lgr5-GFP-DTR* mice (hereafter, Lgr5-GFP) were a kind gift from F. de Sauvage (Genentech, San Francisco, CA)^33^. *Clu^tdTomato^* was generated by crossing *Clu-Cre-ERT2* mice with *Rosa26-CAG-lox-STOP-lox-tdTomato-WPRE* (Jackson Laboratory). *Bac-Clu-GFP* transgenic mice (*Tg(Clu-EGFP)OD95Gsat,,* referred to as *Clu-GFP*) mice were obtained from Rockefeller University (GENSAT project). *Tgfbr2^flox/flox^* (B6;129-Tgfbr2^tm1Karl^) mice were provided by Stephan Karlson (RISE Research Institute of Sweden). *Villin-CreERT2* model was a gift from S. Robine (Institute Curie-CNRS). *Vil-CreERT2; Clu-GFP; Tgfbr2^−/−^* mice were generated in-house by crossing *Tgfbr2^flox/flox^* with *Tg(Clu-EGFP)OD95Gsat* mice and subsequently crossed with *Vil-CreERT2* mice. *Yap^−/−^;Taz^−/−^* mice were generated by crossing Yap*^flox/flox^* or Taz*^flox/flox^* mice with the *Vil-CreERT2* line as previously described^4^. These mice were then crossed with *Tgfbr2^flox/flox^* to generate the *Yap^−/−^;Taz^−/−^;Tgfbr2^−/−^* triple knock out model.

### METHODS DETAIL

#### Isolation of small intestine crypts for multiomics

Crypts were isolated from for multiomics analysis as described in Ayyaz *et al.*^4^. Specifically, small intestines were flushed with ice-cold phosphate buffered saline (PBS), cut longitudinally into roughly 2-mm-long pieces, rinsed with ice-cold PBS, and incubated in 2 mM EDTA at 4°C for 40 min with continuous shaking. The whole-epithelial segments were isolated by vigorous shaking in ice-cold PBS, and enriched for crypts by passing dissociated epithelial segments through a 70 µm mesh cell strainer.

#### Multiomics sample preparation

Nuclei were extracted from isolated crypts following the demonstrated protocol by 10x Genomics (CG000365 revC). Crypts were resuspended in 100 µl chilled lysis buffer (10 mM Tris-HCl pH 7.4, 10 mM NaCl, 3 mM MgCl2, 0.1% Tween-20, 0.1% Nonident P40 Substitute, 0.01% Digitonin, 1% BSA, 1 mM DTT, 1 U/µL RNase inhibitor) and incubated for 5 minutes. 1 mL chilled Wash Buffer (10 mM Tris-HCl pH 7.4, 10 mM NaCl, 3 mM MgCl_2_, 1% BSA, 0.1% Tween-20, 1 mM DTT, 1 U/µL RNase inhibitor) was then added and mixed 5 times. Lysed cells were centrifuged at 500rcf for 5min at 4°C. Nuclei were washed with 1 mL Wash Buffer and centrifuged 2 more times. Nuclei were resuspended in Diluted Nuclei Buffer to approximately 5000 nuclei/µL. Nuclei were then immediately processed with the 10x Genomics Multiome ATAC + Gene Expression kit (PN-1000285) following the commercial protocol (Rev F) for a targeted 10,000 nuclei per sample. Resulting libraries were sequenced on NovaSeq6000 S1 flow cells.

#### Mouse intestinal organoids

Organoids were isolated and cultured according to the protocol established by Sato and Clevers^51^. Thus, crypts were harvested removing the small intestine and cutting it into 3 sections which were then cut longitudinally. These sections were incubated in ice-cold PBS containing 2 mM EDTA at 4 °C for 30 min with slight agitation. The epithelium was separated by vigorous shaking in PBS, and crypts filtered using a 70 μm cell strainer. The resulting crypts were centrifuged for 5 min at 1000 rpm. Crypts were seeded in 24 well plates, each well containing a 35 µL droplet of growth factor reduced Matrigel (VWR, #CACB356231) and grown in 500 µL of media consisting of Advanced DMEM/F12 (Life Technologies, #12634028) supplemented with 2 mM GlutaMax (Life Technologies, #35050061), 100 U/mL Penicillin/100 mg/mL Streptomycin (Life Technologies, #15140122), N2 Supplement (Life Technologies, #17502001), B-27 Supplement (Life Technologies, #17504001), 1.25 mM *N*-acetyl-L-cysteine (Sigma-Aldrich), 50 ng/µl mouse recombinant EGF (Life Technologies, #PMG8043), 100 ng/µl mouse recombinant Noggin (Peprotech #250-38) and either 200 ng/mL human R-spondin1(R&D Systems, #4645-RS) or in conditioned media prepared within the lab and used at 2% after titration. Organoids were maintained by passaging every 4-5 days, which consisted of physical dissociation of Matrigel using an 18G needle and syringe, followed by replating in fresh madrigal and media. Additional culture components included: TGFβ1 (R&D Systems #7754-BH), SB431542 (Selleck Chemicals, #S1067), 5-Florouracil (5-FU; Sigma-Aldrich, #F6627-5G), HTH-02-006 (provided by OICR) and 4-hydroxytamoxifen (4-OHT; Sigma-Aldrich, #H6278; 5 μg/mL). These components were added to culture conditions for various amounts of time based on experimental design, the specifics of which can be found in Figure legends.

#### Lineage tracing experiments (organoids)

Irradiation consisted of a single dose of 5 Gy using a GammaCell 40 irradiator to induce injury 16 hours before passage. To induce Cre-mediated knock out in the organoids (genotype: Clu-^CreERT2+/−^;Rosa26-^LoxP-STOP-LoxP-tdTomato+/+-^;Lgr5-GFP-DTR organoids), 4-OHT was added to culture medium at a concentration of 5 μg/mL 16 hours prior to organoids passaging.

#### Quantification of organoid regeneration

Quantification of lineage traced crypts was performed manually, counting the number of Lgr5-GFP+ regions that displayed ≥ 70% td-Tomato+ signal. To calculate p values, we performed a 2 tailed unpaired T test.

#### Conditional gene knock out and irradiation

Inducible Cre-mediated gene knock-out was performed by intraperitoneal injection of eight- to twelve-week-old mice with two doses of 2 mg Tamoxifen (Sigma-Aldrich, #15648-1G) in corn oil, administered consecutively. Both sexes of mice were used for experimental procedures. Epithelial injury was achieved by exposure to single dose of 12 Gy irradiation using a GammaCell 40 irradiator. Mice were euthanized by CO_2_ asphyxiation. The number of animals used for each experiment is indicated in the figure legend as *n=x* where *x* represents the number of animals per group. All experiments were performed according to approved procedures by the Canadian Council on Animal Care (CCAC) and following the ethical guidelines for responsible use of laboratory animals in biological research, taking a reductionist approach. All procedures were reviewed and approved by the Animal Care Committee (ACC) at the Lunenfeld-Tanenbaum Research Institute (LTRI).

#### Human colon organoid culture

Organoids were cultured according to the guideline given by the Princess Margaret Living Biobank (PMLB), where they were primarily established following colon tissue biopsy. Organoids were seeded in 24 well plates, each well containing a 35 µL droplet of growth factor reduced Matrigel (VWR, #CACB356231) and grown in 500 µL of media consisting of Advanced DMEM/F12 (Life Technologies, #12634028) supplemented with 2 mM GlutaMax (Life Technologies #35050061), 100 U/mL Penicillin/100 mg/mL Streptomycin (Life Technologies, #15140122), B-27 Supplement (Life Technologies, #17504001), 50 ng/µL mouse recombinant EGF (Life Technologies, #PMG8043), 100 ng/µL mouse recombinant Noggin (Peprotech, #250-38), 1.25 mM *N*-acetyl-L-cysteine (Sigma-Aldrich, #A9165), 10 µM SB20219010 (Sigma-Aldrich, #559388), 10 nM Gastrin I (Sigma-Aldrich, #G9145), and 0.5 µM A83-01 (Selleck Chemicals, #S7692). Additional media components, WNT3a and RSPO-1 conditioned media, was provided by the PMLB. Organoids were maintained by passaging every 4-5 days, which consisted of a10 minute incubation in TryplE (ThermoFisher Scientific, #12604013) at 37°C overnight, followed by physical dissociation using an 18G needle and syringe, and replating in fresh matrigel and media.

#### Validation of gene knock out in Tgfbr2^flox/flox^ mice

The small intestines were isolated as described above from *Tgfbr2flox/flox* x *VilCre* mice that were either un-injected or injected with Tamoxifen for 24 or 48 hours. As a negative control, an un-injected WT mouse was also used. Epithelia was isolated and incubated in Trypsin-0.25% EDTA to make single cells. Epithelial cells were selected out using EpCAM antibody (Biolegend #18206) and magnetic beads. DNA was extracted from the cells and 60 ng of DNA was used for PCR to detect WT - 422 bp, floxed - 575 bp (P3: 5′-TATGGACTGGCTGCTTTTGTATTC-3′ x P4: 5′-TGGGGATAGAGGTAGAAAGACATA-3′) and excised/null – 695 bp (P3: 5′-TATGGACTGGCTGCTTTTGTATTC-3′ x P5: 5′-TATTGGGTGTGGTTGT GGACTTTA-3′) TβRII gene. PCR conditions are as follows 94 C° for 2 min; 10 cycles (94 C° for 20 sec, 65 C° for 15 sec, 68 C° for 10 sec) followed by 25 cycles (94 C° for 15 sec, 60 C° for 15 sec and 72 C° for 10 sec); 72 C° for 2 min.

#### Tissue preparation

Unirradiated and irradiated mice were sacrificed and the small and large intestines were extracted. The small intestine was cut into 3 equal-length sections representing the duodenum, jejunum and ileum, and each section was flushed with cold PBS. Intestines were prepared using 2 methods. Sections were cut into approximately 1cm segments and separated by section into cassettes before being placed overnight in 10% Neutral buffered formalin (NBF; Fisher Scientific, #SF100-4) stored overnight at room temperature, with shaking. Alternatively, Swiss rolls were prepared by flushing the sections with modified Bouin’s fixative (50% ethanol, 5% glacial acetic acid) cut longitudinally, and rolled luminal side up from the proximal to distal end of the section. These were fixed overnight in 10% NBF. Samples prepared for cryopreservation were transferred to 30% sucrose in PBS, and stored overnight at 4°C with shaking, followed by embedding and freezing in OCT (VWR, #95057-838). Alternatively, tissue in a cassette was prepared for paraffin embedding by dehydration in an ethanol series (25%-5 min; 50%-5 min; 70% rinse; 70%-10 min; 100%-30 min-repeat 3 times) and incubation in Histo-Clear II (1 Hr, repeat 3 times)(Diamed, #NDIHS202). The tissue cassettes were transferred to a container of paraffin (VWR, #15159-464) for 15 minutes, followed by overnight incubation in fresh paraffin at 60°C.

#### Immunohistochemistry

Paraffin-embedded tissue sections were deparaffinized by incubating twice in Histo-Clear II solution for 15min each and rehydrated for 2mins each in 100% (2x), 95% and 70% ethanol respectively. Slides were then boiled at ~95°C in 10 mM Sodium Citrate (Sigma-Aldrich, #S4641-1KG) buffer supplemented with 0.05% Tween-20 (Sigma-Aldrich, #P9416) for 15 mins and cooled on ice for 10 mins followed by washing in 1X PBS (Multicell, #311-425-CL) for 5 mins. Subsequently, slides were blocked for 1 hour at room temp. in 1X PBS supplemented with 5% bovine serum albumin (BSA) (Sigma-Aldrich, #A7906) and 0.3% Triton X-100 (Sigma-Aldrich, #X100). After which the slides were incubated overnight at 4°C with primary antibody diluted in 1X PBS supplemented with 1% BSA and 0.3% Triton X-100. Next day, slides were washed twice in 1X PBS for 5mins and incubated with Dako EnVision+ System-HRP (Dako, #K4003) for 1 hour at room temp. Followed by two washes in 1X PBS for 5 mins and then incubated with 3,3′-Diaminobenzidine (DAB) Liquid Substrate (Sigma-Aldrich, #D6190) for 5 mins at room temp. Slides were again washed twice in 1X PBS for 5 min each and counterstained with Harris modified Hematoxylin solution (Sigma-Aldrich, #HHS16) for 30 seconds followed by rinsing in tap water for 5 mins and quick de-staining in acid ethanol (95% ethanol and 5% HCl). Slides were then rinsed in tap water for 5 mins and dehydrated for 2 mins each in ethanol at 95%, 100% and Histo-Clear II respectively. Slides were then mounted with Histomount medium (National Diagnostics, #HS-103) and dried overnight at room temp. Images were acquired at 20X magnification using a ZEISS Axio Scan.Z1 Slide Scanner. Antibodies: Rabbit anti-Yap1 (Cat# 14074S; 1:300) and Rabbit anti-Smad2/3 (NEB #8685S; 1:300).

#### RNAscope in situ hybridization

smFISH (RNAScope; Advanced Cell Diagnostics) was performed on paraffin embedded sections according to the manufacturer’s recommendations, using the RNAscope Multiplex Fluorescent Reagent Kit v2 Assay. Tyramide signal amplification (TSA) fluorophores (Akoya Bioscience, TSA Fluorescein #NEL741001KT, TSA Cyanine 3 #NEL744001KT, TSA Cyanine 5 #NEL745E001KT) were used for detection. The following probes were acquired from ACD: mm-Clu-C2(#427891-C2), mm-Lgr5-C3(#312171-C3), mm-ki67-C1(#416771), mm-Cde3-C3 (#314721-C3), mm-Pdgfra-C4 (#480661-C4), mm-Adreg-C3 (#460651-C3) and mm-Tgfbr2 (#406241). Quantification and analysis of RNAScope was performed using Cell Profiler software.

#### Fluorescent in situ hybridization using RNA probe

For synthesizing fluorescein-labelled *Smad7* antisense RNA probe, a plasmid with 0.6kb EcoRI-SalI fragment of mouse *Smad7* cDNA coding region cloned into pBluescript KS- was used. *In vitro* transcription was performed using Fluorescein RNA Labeling Mix (Millipore Sigma, #11685619910). Paraffin sections were deparaffinized and rehydrated using standard procedures, then incubated with 3% H_2_O_2_ in PBS with 0.1% Tween20 (PBS-T) for 30 min at RT to quench endogenous peroxidase activity. The sections were incubated in 10 mM Tris-HCl, 1 mM EDTA, pH 9.0 at 95C for 20 min, followed by digestion with 1 μg/mL proteinase K in PBS-T at 37°C for 30 min and post fixation using 4% paraformaldehyde in PBS at RT for 10 min. After pre-hybridization for 1h at 65° C, the slides were incubated with probe diluted in hybridization buffer at 65 °C overnight. The slides were washed twice in 1x SSC, 50% formamide at 37 °C for 30 min each. Blocking was performed using 2% Blocking Reagent (Millipore Sigma, #11096176001) in PBS-T, prior to antibody incubation to detect the probe. Hybridized probe was detected using peroxidase conjugated anti-fluorescein antibody (Millipore Sigma, #11426346910, 1:1000) and TSA Vivid 520 fluorophore kit (Bio Techne, #7523). DAPI (0.5 μg/mL) was used for nuclear counterstain.

#### Hematoxylin and eosin (H&E) staining

Paraffin-embedded tissue sections were deparaffinized by incubating twice in Histo-Clear II solution for 15 min each. Followed by rehydration in 100% (2x), 95%, 70% ethanol and double distilled water (ddH2O) for 2 mins respectively. Slides were then stained with Harris modified Hematoxylin solution (Sigma-Aldrich, #HHS16) for 30 seconds and rinsed in tap water for 5mins. Followed by quick de-staining in acid ethanol (95% ethanol and 5% HCl) and further rinsing in tap water for 5mins. Slides were then counterstained with 0.25% Eosin Y solution (Sigma-Aldrich, #HT110116) for 5 seconds and quickly rinsed in ddH2O before dehydration with ethanol at 70%, 95%, 100% and Histo-Clear II solution for 1min each respectively. Slides were then mounted with Histomount medium (National Diagnostics, #HS-103) and dried overnight at room temp. Images were captured at 20X magnification using a ZEISS Axio Scan.Z1 Slide Scanner.

#### Immunofluorescence

##### Intestinal sections

Cryosections (16-μm thick) permeabilized in PBS with 0.5% Triton-X-100 (Sigma-Aldrich, #X100-500ML) and 0.2% Tween-20 (Sigma-Aldrich, #P1379) for 30 min, and blocked in 2% BSA in PBS for 1 h. Sections were incubated with primary antibodies, diluted in PBS with 0.2% Tween-20 and 0.5% BSA (Sigma-Aldrich, #A7906) at 4 °C overnight. Paraffin sections (5-μm thick) were prepared in the same manner as indicated in IHC staining above.

##### Organoids

Organoids were grown on glass coverslips after passaging. At the desired timepoint, organoids were fixed in 4% paraformaldehyde (Electron Microscopy Sciences, #15710) in PBS for 30 minutes, followed by permeabilization and blocking for 1 hour at room temperature in PBS with 0.5% Triton-X-100, 0.2% Tween-20 and 2% BSA. Coverslips were incubated overnight at 4°C in primary antibodies diluted in PBS with 0.2% Tween-20 and 0.5% BSA.

The following primary antibodies were used: rabbit anti-lysozyme (Dako, #2230, 1:1,000), rabbit anti-Muc2 (Santa Cruz, #sc-15334, 1:300), rabbit anti-Olfm4 (Cell Signaling, #66479, 1:300), rabbit anti-ChgA (Abcam, #ab85554, 1:250), goat anti-GFP (Abcam, #ab5450, 1:1000), chicken anti-GFP (Abcam, #ab13970; 1:1000) Rabbit anti-ki67(Abcam, #ab15580), Anti-CD326 (Biolegend, #118205), Rabbit anti-Smad2/3 (Cell Signaling, #8685S, 1:300), Rabbit anti-pSmad2 (Cell signaling, #18338, 1:200), Rabbit anti-Smad7 (Abcam, #ab216428; 1:100). Subsequently, sections and coverslips were incubated with DyLight Fluor secondary antibodies (Fisher Scientific) diluted in PBS with 0.2% Tween-20 and 0.5% BSA overnight at 4°C. Nuclear counterstain was done with with 0.1 µg/mL DAPI (Sigma-Aldrich, D9542) for 30 min before mounting with a mounting medium based on Mowiol 4-88 (Sigma-Aldrich, #81381). The slides were then left to dry overnight at room temperature. Images were acquired using either a Nikon A1RHD or AXR confocal microscope platform and analyzed using Nikon NIS-Elements HC and ImageJ software.

#### Flow cytometry

Single cells were prepared from organoids cultures by incubating organoids in Trypsin (VWR #CA45000-664) for 10 minutes followed by mechanical dissociation via 18G needle and 12 mL syringe. Cells were diluted in Advanced DMEM F12 (Life Technologies #12634028) and spun at 1000 rpm prior to being resuspended in FACS medium (ice-cold PBS containing 25 mM HEPES (Gibco) and 1% fetal bovine serum (VWR). No antibody staining was performed, and cells were sorted using endogenous GFP signal. DAPI (0.1 µg/mL) was added to the final FACS medium for 30 min before flow cytometry analyses, which was performed using the BD Fortessa X-20 benchtop flow cytometer. Non-GFP-expressing organoids were used to provide a negative control for endogenous Clu-GFP. Analysis was performed using Kaluza Software.

#### Fluorescence Activated Cell Sorting

Single cells were prepared from organoids cultures by incubating organoids in Trypsin (VWR, #CA45000-664) for 5 minutes at 37°C on a shaker. After 5 minutes, cells were pipetted vigorously and placed back at 37°C, with process repeated until organoids were sufficiently dissociated to single cells. Cells were diluted in Advanced DMEM F12 (Life Technologies, #12634028) supplemented with 10% FBS and spun at 1000 rpm prior to being resuspended in 10 mL ice cold PBS. Following aspiration of PBS, cells were resuspended in 500 µL FACS medium (ice-cold PBS containing 25 mM HEPES (Gibco) and 1% heat inactivated fetal bovine serum (VWR) and DAPI (0.1 µg/mL)) and strained through a 40 µm cell strainer. Sorting was performed by Lunenfeld-Tanenbaum Research Institute technician using the Astrios EQ Cell Sorter. GFP-/tdTomato-organoids were used to provide a negative control for endogenous fluorescent proteins. Additional tdTomato+ organoids (*Clu-cre-ERT2;Rosa26-CAG-lox-STOP-lox-tdTomato)* were used as gating controls.

#### Organoid propagation from single cells

Following collection, organoids were plated in Matrigel in 24 well plates. Cells were diluted to 10, 000, 5, 000, 1,000 and 500 cell concentration in 35 µL droplets of Matrigel. They were grown for 2 days in ENR media containing CHIR-99021 (Sigma-Aldrich, #252917-06-9; 3 μM) and Y-27632 (Stem Cell Technologies, #72307; 10 μM). Media was replaced with normal growth (ENR) media, that was changed every 2-3 days until organoids formed crypts, up to d14 after sorting.

#### Quantitative real-time PCR

A RNeasy Mini Kit (Qiagen) was used to extract total RNA. Superscript IV Reverse Transcriptase (Life Technologies, #18090200) was used to synthesize cDNA according to the manufacturer’s instructions. We used oligo(dT) primer and 0.1-0.8 µg of RNA per synthesis reaction. cDNA was mixed with forward and reverse primers (0.5 mM each), nuclease free water and SYBR Green I Master mix (Roche, #04707516001) to a final volume of 15 µL/ well in a 384 well plate, with each sample run in triplicate. Detection was achieved using a Roche Light-Cycler, with conditions set for a 10 min of initial incubation at 95 °C and 42 cycles of 95 °C, 60 °C and 72 °C (10 seconds/temperature) for amplification. expression was calculated using the ΔΔCt method and normalized to *Ywhaz*.

#### Multiome data preprocessing

To begin our multiomic analysis, we first created a mouse genome using the ensemble mm39 dataset following 10x Genomics’ recommendations. Then, using the default settings for CellRanger ARC 2.0, we processed the sequences from the 0 and 2 dpi samples. The raw output was then analysed using the Seurat/Signac pipeline as follows^16,52^. First, CellRanger detected peaks from both datasets were read, common and overlapping peaks were merged, and all peaks were combined to create one global peak set for both time points. The global peakset was then filtered to remove widths of more than 10000 bp and less than 20 bp. The RNA outputs were then read in using the Read10X function and these were converted into Seurat objects using the CellRanger metadata outputs for use in quality control filtering^16,52^. A counts matrix for the global peakset was constructed for each timepoint using the respective fragment files and gtf file of the reference genome to provide annotations; the counts matrices were then added to the Seurat objects made previously.

To obtain further quality control metrics for the ATAC data, we calculated the nucleosomal signal, transcription start site (TSS) enrichment, the percent of reads in peaks, and the fraction of reads in blacklist regions. For the RNA data, we also calculated the percentage of mitochondrial reads per nuclei. Because we were using an updated version of the mouse genome, we also updated the blacklist regions with Liftover (https://genome.ucsc.edu/cgi-bin/hgLiftOver). After filtering out genes detected in less than 3 nuclei, we filtered low quality nuclei with greater than 50% mitochondrial reads, 5% reads in blacklist regions, nucleosomal signal of 4, and nuclei with less than 250 unique molecular identifiers (UMIs) and genes, 100 detected peaks, percent of reads in peaks of < 15%; for dataset specific filters we selected for nuclei greater than 2.0 and 2.25 TSS for 0 dpi and 2 dpi datasets. Then, using the scater and scran packages, we performed normalization as we did previously^4^ and filtered out doublets using DoubletFinder^53^.

For the ATAC assays for both datasets we performed term frequency-inverse document frequency (TFIDF) normalization, selected top features, and performed singular value decomposition (SVD). We then merged both datasets into one object and next embedded cells. For use with downstream analysis, we also created a custom data package using the BSgenome packaged in R following the developers’ recommendations. To embed cells after merging, we considered each modality on its own and then both. For RNA based embedding, we selected the top 2000 variable genes, scaled, performed PCA and selected the first 20 as most informative of the underlying topology of the data with an elbow plot. Using these 20 principal components, we embed the cells using UMAP. To cluster cells, we performed K-means clustering at increasing resolutions from 0.1 to 2.0 at increments of 0.1; using the clustree package, we selected a resolution of 1.0 as it best-separated cells based on expected celltypes and lowest level where the clusters did not break down into further clusters at the next higher resolution. To embed cells using the ATAC assay, we first recalled peaks using MACS2, kept peaks from standard chromosomes using the keepStandardChromosomes function, and then retained peaks that were not in the blacklist regions. Similar to how the peaks count matrices were formed for the individual datasets, a count matrix for the new recalled peaks was constructed and added to the Seurat object. To aid in convenience of coverage analysis, we also ‘linked’ each gene to its corresponding peak by first computing the GC cotent per peak using the RegionStats function with our custom BSgenome package followed by the LinkPeaks function for all genes in the RNA assay. Using this count matrix, we then performed TFIDF normalization, found the top features, performed SVD, and then ran UMAP using the second (2) to thirty (30) dimensions; we then performed K-means clustering using these dimensions and used clustree as previously described cluster peaks-based embedding of cells at a resolution of 0.5. For multi-modal embedding, we made use of Seurat’s weighted nearest neighbour approach^16^. First, we calculate multi-modal neighbors using the first 20 principal components calculated for the RNA assay and the 2:30 dimensions calculated for the ATAC assay. Using this graph, we performed UMAP as well as K-means clustering as we did for RNA and ATAC to settle on a clustering resolution of 0.5.

For the RNA and the WNN embedded cells, we used cell type specific markers to assign cell identities to clusters. For both cases, we noticed that because there were so few Tuft Cells, we manually picked the cells using the CellSelector function and assigned the identity to Dclk+ and Trpm+ cells^54^. Once individual clusters had been assigned to a celltype, we then collapsed multiple clusters belonging to the same celltype as agglomerative clusters (i.e., multiple EC clusters were collapsed into a general EC cluster). To assign general identities to the ATAC embedded cells, we calculated the percent overlap between each ATAC cluster and each WNN cluster using the metadata information from both types of clustering. Using the gross celltype identities, we then performed differential peak analysis for positively enriched peaks across clusters using the FindAllMarkers function set to use the LR differential test and rowMeans function (**Table 3**). Highly enriched peaks for each cell type were then determined using a knee-based method. For this method, peaks obtained from the FindAllMarkers function for each celltype were ranked according to their −log10 p-value and then the ‘knee’ of the rankings was found using the KneeLocator function of the Kneed python package with the curve parameter set to ‘convex’ and the direction set to ‘increasing’. Once the knee was found, we took all peaks from 100 peaks to the left of the knee and up to the highest rank for each cell type. The peaks obtained from all celltypes are then combined together to form one ‘library’ to use for visualization (**Table S4**). To visualize enriched peaks, we took this library and used it to find the cluster level average for each identified peak using the counts slot of the MACS2 calculated peaks matrix in the Seurat object and performed 2D hierarchical clustering using the Seaborn python package and the scipy package, first creating a distance matrix between peaks using the scipy.cluster.hierarchy.linkage function and then scipy.cluster.hierarchy.fcluster and manual polishing to yield 12 distinct clusters that we overlaid onto the 2d clustered heatmap. Finally, for each peak-cluster, we quantified the proportion of each cell type that contributed accessible peaks to the cluster and visualized the results as a stacked bar-plot.

#### Enterocyte Zonation for WNN Embeddings

To categorize the EC clusters in our WNN embeddings as Crypt Villus Junction (CVJ), Villus Bottom, Villus Middle, and Villus Top, we used the zonation gene sets we had previously used in Ayyaz *et al.*^4^, that were derived from the Moor *et al.*^26^ dataset. Then, we took the RNA normalized counts in the data slot of the Seurat object out of log-space by transforming them with the expm1 function to calculate the mean expression per nuclei of each zonation specific geneset. Clusters were categorized as being members of a particular zone based on which zonal signature was highest in the EC clusters.

#### Motif Analysis

To analyze motifs across the dataset, we followed Signac’s instructions for motif analysis. In brief, we downloaded the latest JASPAR2022 dataset of position frequency matrices (PFMs) for vertebrates and then created a Motif class object to add this information to the MACS2 chromatin accessibility matrix of our Seurat object using the AddMotifs function, alongside our custom BSgenome data package. Then, we took the cluster specific enriched peaks for each gross cell type, filtered for those with a p-value < 0.005, and looked for motifs using the FindMotifs function with Signac using the default settings. Under these settings, Signac will use these cluster specific peaks and will perform a hypergeometric test that compares how enriched each motif binding element defined in the JASPAR2022 dataset is compared to the full MACS2 identified peakset of our multiomic dataset. The result of these calculations is a list of cluster specific motif binding elements. Signac does not provide single nuclei level measurements of motif enrichment, thus we opted to use chromVAR to calculate this information. Then we exported the chromVAR matrix for our nuclei and loaded it into python to create a dotplot^9^. We also read in the Signac list of cluster specific motifs and took the top 150 motifs out of all motifs identified per celltype and created a global ‘motif set’. Using this global motif set, we found the average chromVAR Z-score for each individual motif across the nuclei for each cluster to arrive at cluster level chromVAR Z-score averages. As an example, we found the enterocyte level average chromVAR Z-Score of the SMAD2 motif by the averaging the chromVAR Z-scores of the motif across all nuclei identified as enterocytes and repeated this calculation for all remaining motifs. We then used a Seaborn script to visualize these results of select celltype specific motifs as a dotplot with the size of each circle corresponding to the relative percentage of nuclei of a given cluster as being greater than or less than zero depending on whether the cluster level chromVAR Z-score average of a motif was positive or negative respectively. For visualization we capped the color scale on a range of −1 to +1 and then created visualizations for select revSC motifs and epithelium wide motifs. We did not show results for TCs and fibroblasts as there were not many nuclei from these celltypes.

#### Convergence Analysis

To examine the convergence between Hippo and Smad signalling pathways, we made use of the motifMatchr package in R. In brief, we took the PFMs for TEADs 1-4 and SMAD 2/3 from the JASPAR2022 dataset for vertebrates and identified revSC selective peaks (with p-value <0.005). To examine overlap of motifs per peak we used motifMatchr’s matchMotif function in conjunction with the JASPAR2022 PFMs, the revSC selective peaks (p<0.005) as the query, the background motif frequencies of the global peakset, and our custom GCRm39 reference BSGenome package to calculate the per peak overlap between TEADs1-4 and SMAD2/3. For analysis, we categorized all TEAD containing peaks as one category of TEADs. Then we used the vennDiagram package in R to determine the overlap between TEAD containing and SMAD2/3 containing peaks. Then, we performed Fischer’s Exact t-test using the hypergeometric distribution to calculate p-values for the different combinations of overlap of the queried transcription factors, for the numbers of TEAD-only and SMAD-only containing peaks, and finally, the number of selective revSC peaks containing a Tead or SMAD site.

Next, we performed the same analysis for the global peakset to determine how many of the global peaks were TEAD specific, SMAD specific, and SMAD and TEAD specific. We then calculated the mean accessibility for each category (mentioned above) per nuclei and visualized the results using Seurat’s violin plot function. To calculate p-values, we then performed a two tailed unpaired t.test assuming equal variance comparing the mean accessibilities of SMAD, TEAD, and SMAD and TEAD containing peaks between revSCs and other epithelial cell types.

#### Trajectory inference of enterocytes

EC, RevSC, and ICC clusters from both 0dpi and 2dpi samples were first subset. The sample-matched anoikis-like EC-W2 and EC-W7 clusters were omitted from the analysis given their extremely low transcript content and lack of any expression pattern specificity. As such, they likely reflect apoptotic cells and could introduce artifacts in downstream analysis. Of note, while transcript complexity was very low, the quality of the ATAC profiles did not appear compromised. To ensure that these populations didn’t reflect terminal villus populations, which are known to undergo anoikis, we confirmed that they did not express established markers (*Ada*, *Nt5e*). Remaining cells from the two samples were then aligned using scVI (scvi-tools v0.19.0)^24^. UMI counts were used as input, supplying the sample ID as the batch_key. Other parameters include n_layers=2, encode_covariates=T, deeply_inject_covariates=False, use_layer_norm=”both”, use_batch_norm=”none”. Model training was performed with max_epochs=400, early_stopping=True, early_stopping_monitor=”elbo_validation”, and early_stopping_patience=10. The model converged after 115 epochs.

The 10-dimensional latent representation was then used to re-embed and re-cluster the data using scanpy’s pp.neighbors, tl.umap, and tl.leiden (resolution=0.2) functions with default parameters. This resolved two rare clusters of cells not involved in the dominant EC developmental continuum: one expressed goblet cell markers (observable on the UMAP embedding of the whole dataset), while the other had expression patterns consistent with colonocytes/distal ileal enterocytes (*Fabp6*, *Naaladl1, Slc51a*)^40^. These clusters were under-represented and were distinct from the dominant developmental trajectory, were removed, and the data re-embedded using the latent representation from scVI and re-clustered at a resolution of 0.1. The resulting embedding is shown in the main figures and reflects phenotypes spanning from Lgr5+ ISCs to Ada+ villus tip enterocytes.

To project the cells onto a continuous trajectory spanning these phenotypes, the resulting UMAP embedding was provided as input for slingshot (v2.4.0)^25^ with start.clus=1 (the LGR5+ cluster) and thresh=1. Resulting pseudotime values were scaled to range from 0-1. Discrete states along the continuum were defined by segmenting the cells into 5 equally sized bins by pseudotime value.

#### Regional differential expression and chromatin accessibility

Both differential expression and chromatin accessibility were evaluated to assess homeostatic zonation patterns (i.e. as a function of a 0dpi cell’s position along the inferred trajectory), and the effects of irradiation on phenotypes along the CV axis (i.e. 2 dpi and 0 dpi cells mapped to the same region on the trajectory). Differential expression was performed fitting a negative binomial generalized additive model (NB-GAM) as implemented by the R package tradeSeq (v1.10.0)^55^ with nknots=6, and testing for an association with pseudotime value using the package’s associationTest() function.

Differential chromatin accessibility was evaluated using the logistic regression model implemented in Seurat’s (v4.3.0) FindMarkers() function^16^, including the log10-transformed raw read count per cell as a covariate. We opted to not include ATAC fragment counts as a covariate due to the biologically relevant changes in total accessibility throughout differentiation. Differentially accessible peaks (adjusted p < 0.05; logFC > 0.5; min.pct = 0.01) were evaluated for transcription factor motif over-representation using the hypergeometric test implement by Signac’s FindMotifs() function with default parameters and the JASPAR2022 collection of motifs.

#### Zonation marker gene set scoring

Sets of genes with distinct zonation patterns were collected from Moor *et al*.^26^. Seurat’s AddModuleScore was used to calculate gene set activity scores for each cell using default parameters. Expression patterns of the gene sets were then examined across the inferred developmental trajectory.

### QUANTIFICATION AND STATISTICAL ANALYSIS

Statistical details of experiments can be found in the figure legends and main text above. Analysis was performed using R, Python, and Graphpad Prism. For all experiments, significance was determined with α < 0.05. When appropriate, multiple hypothesis testing correction was performed using the Benjamini-Hochberg procedure at a false discovery rate <0.05.

## Supplemental Information

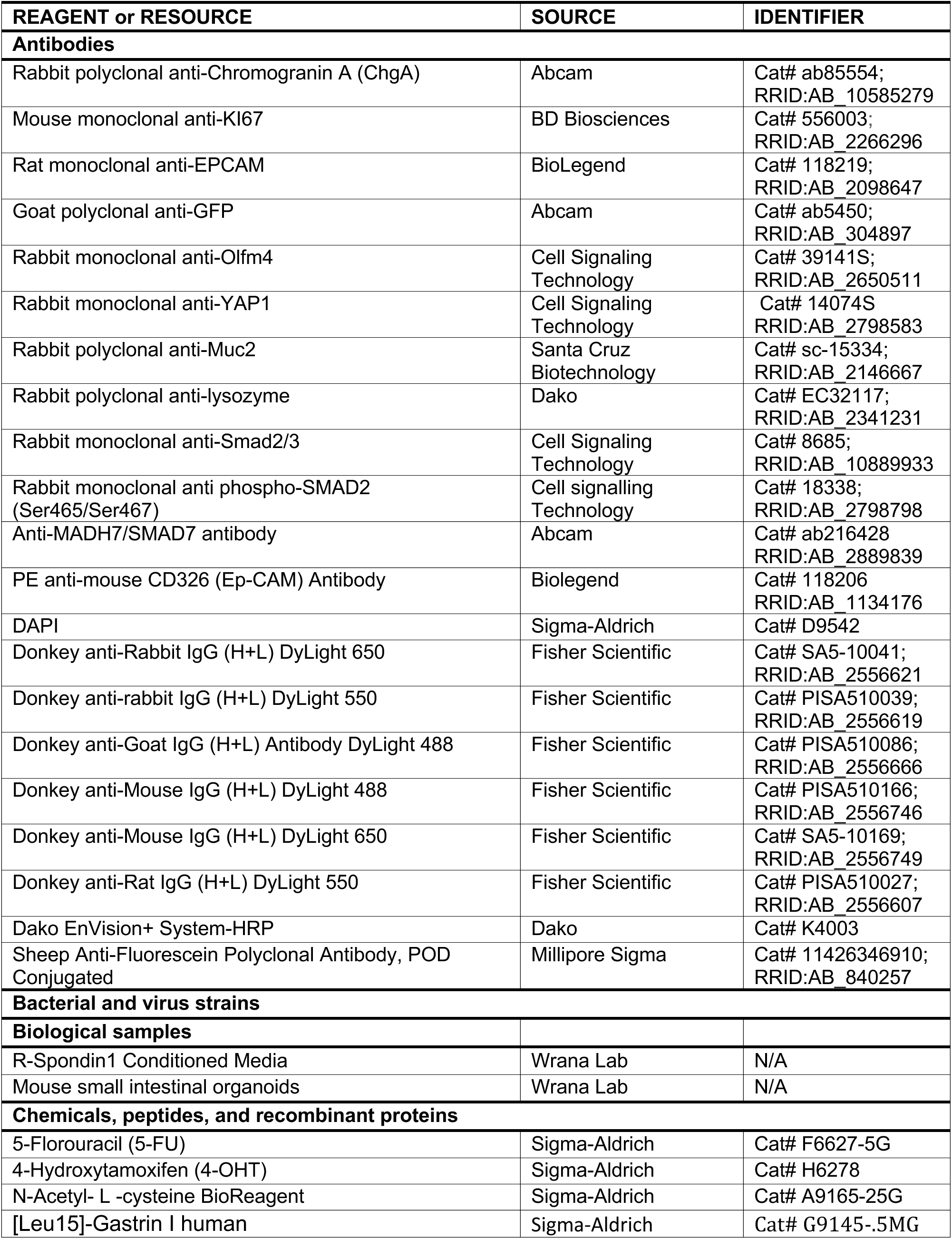
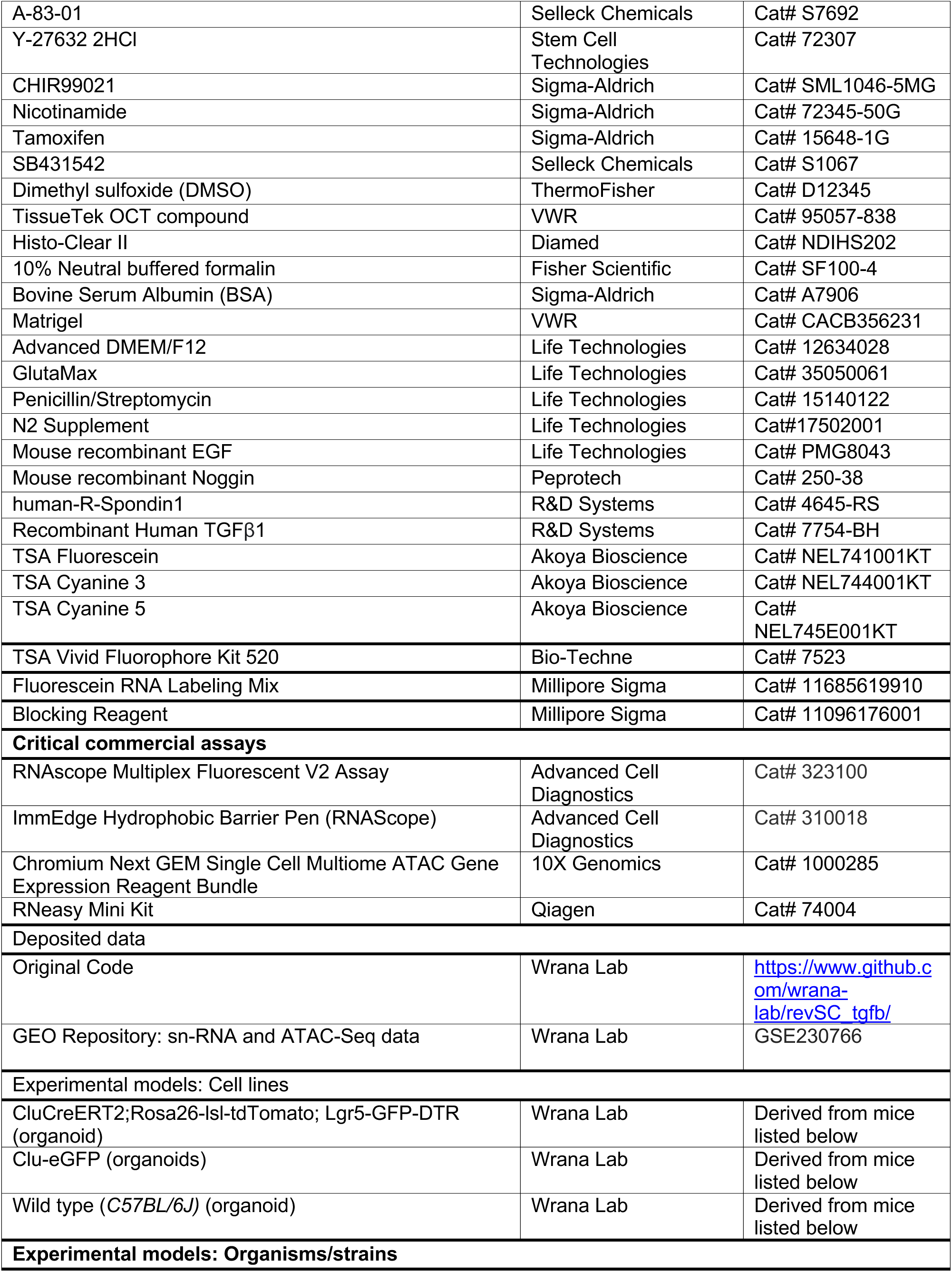
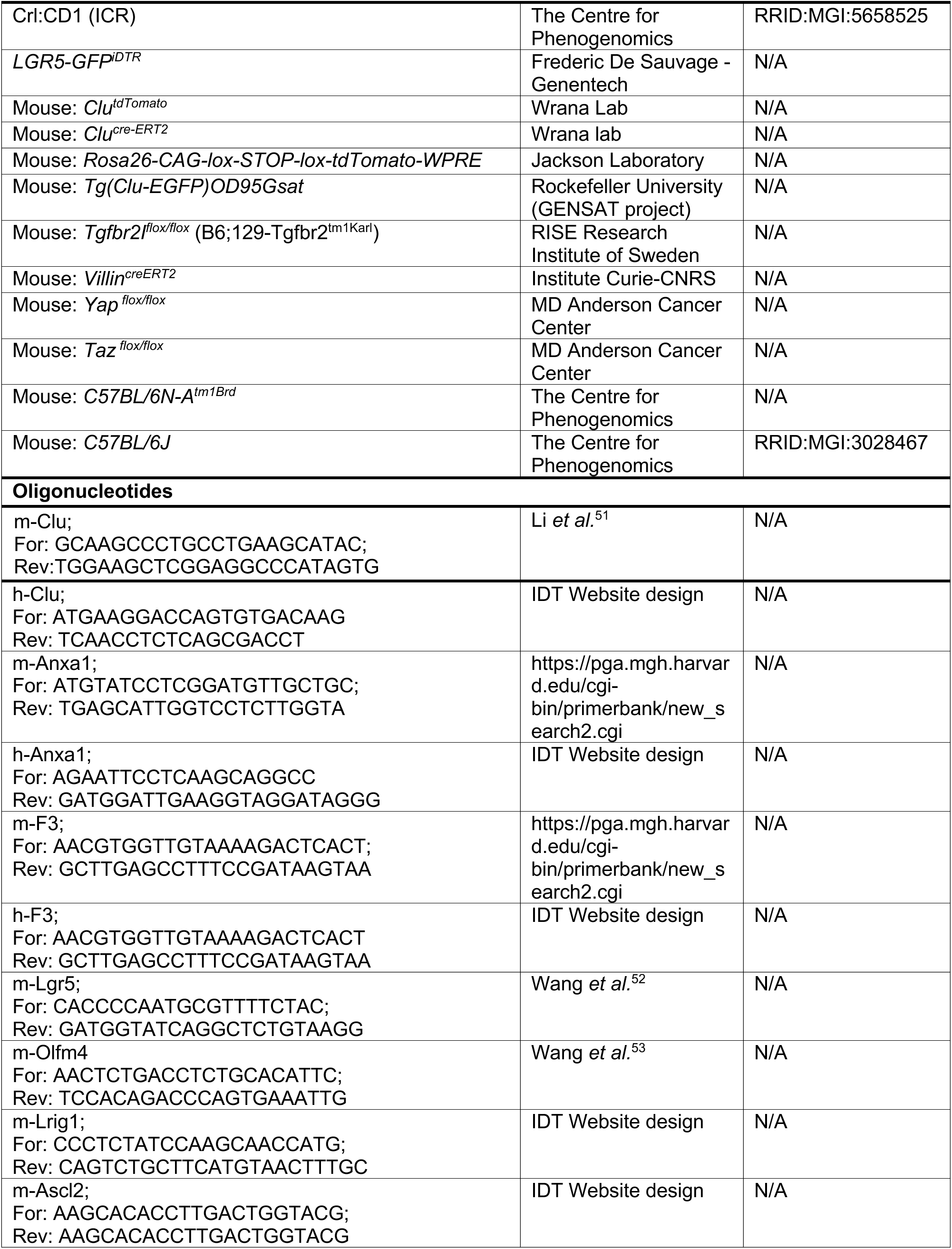
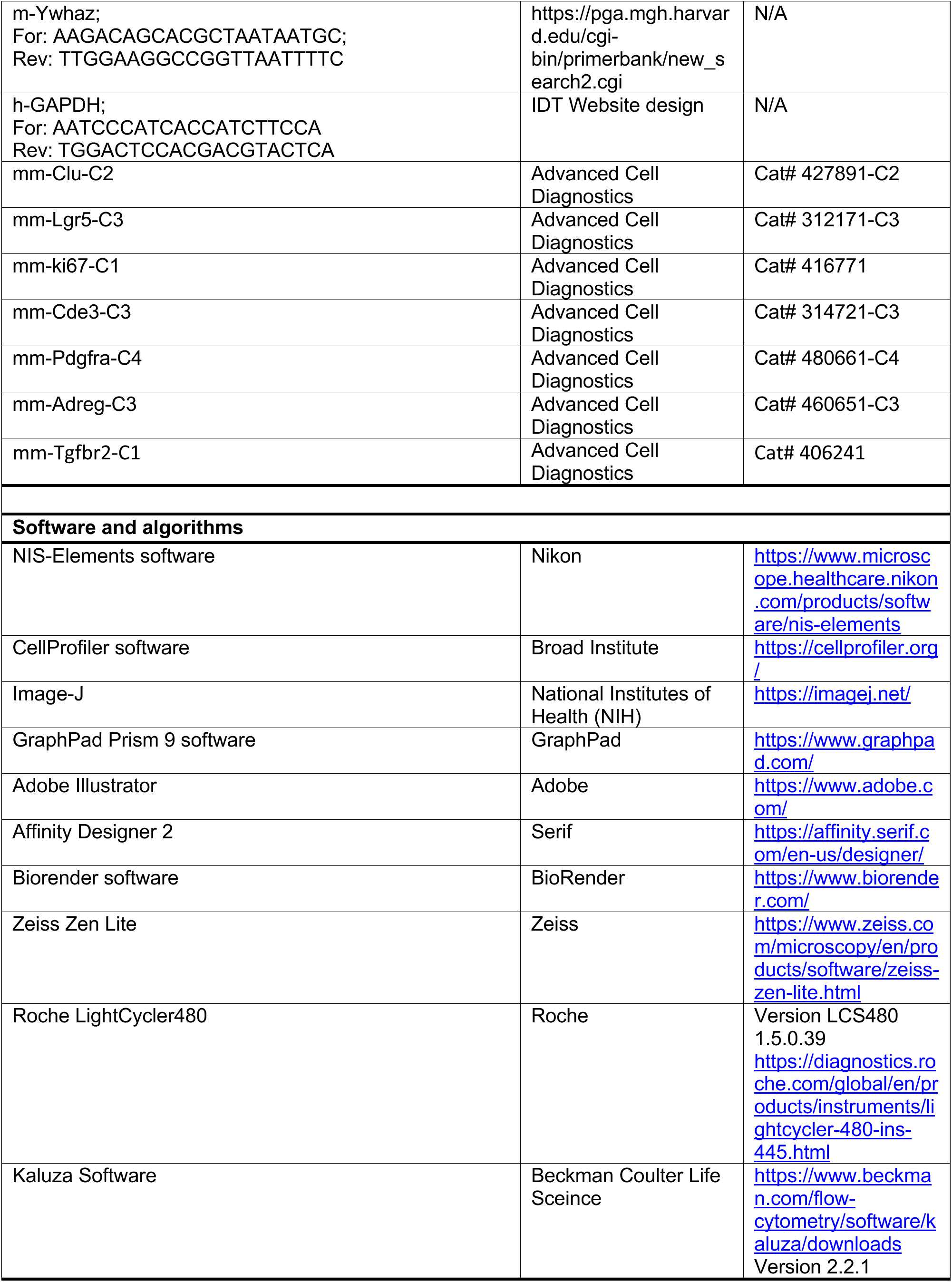
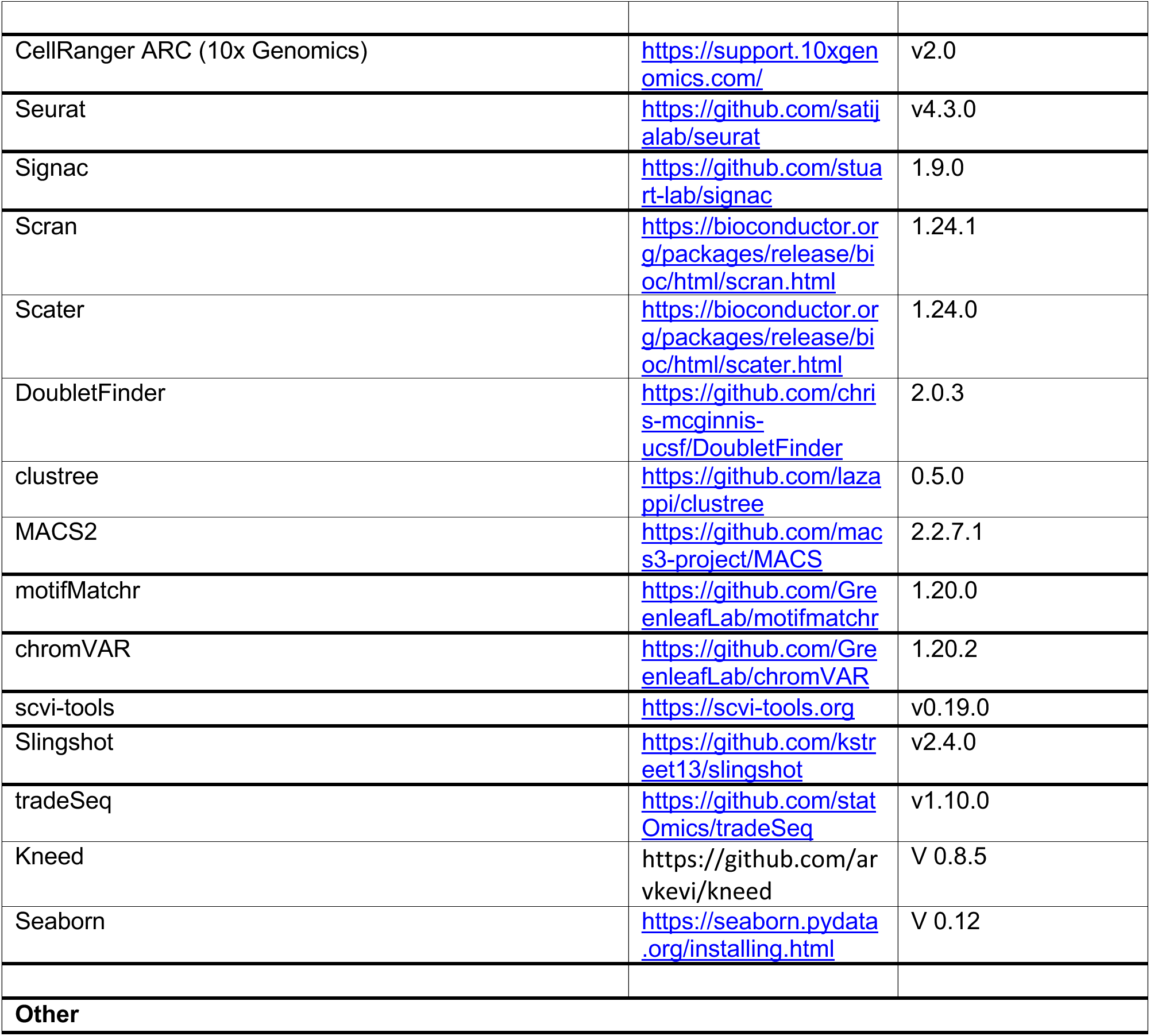

## Extended Data and Figure Legends

### Extended Data Table Legends

**Table S1.** Differentially expressed genes associated with transcriptome clusters in ED Fig. 1c

**Table S2.** Differentially enriched peaks associated with ATAC-Seq clusters in Figure 1

**Table S3:** ATAC peaks associated with major intestinal lineages

**Table S4:** Differentially enriched motifs in epithelial cell types

**Table S5**: ATAC peaks highly enriched in distinct epithelial lineages

**Table S6**: ATAC peaks from Table S5 accessible in different lineages

**Table S7.** Transcription factor motifs enriched in zonation-associated regulatory elements

**Table S8.** 50-gene signature of revSCs

**Table S9.** Differentially expressed genes along the crypt-villus axis at 2 dpi

**Table S10.** Transcription factor motifs enriched in 2dpi-specific peaks along the crypt-villus axis

**Table S11**: RevSC selective peaks containing SMAD only, TEAD only, or SMAD and TEAD sites

**Table S12:** Global peak set of SMAD only, TEAD only, or SMAD and TEAD Sites

**Extended Data Fig. 1.**
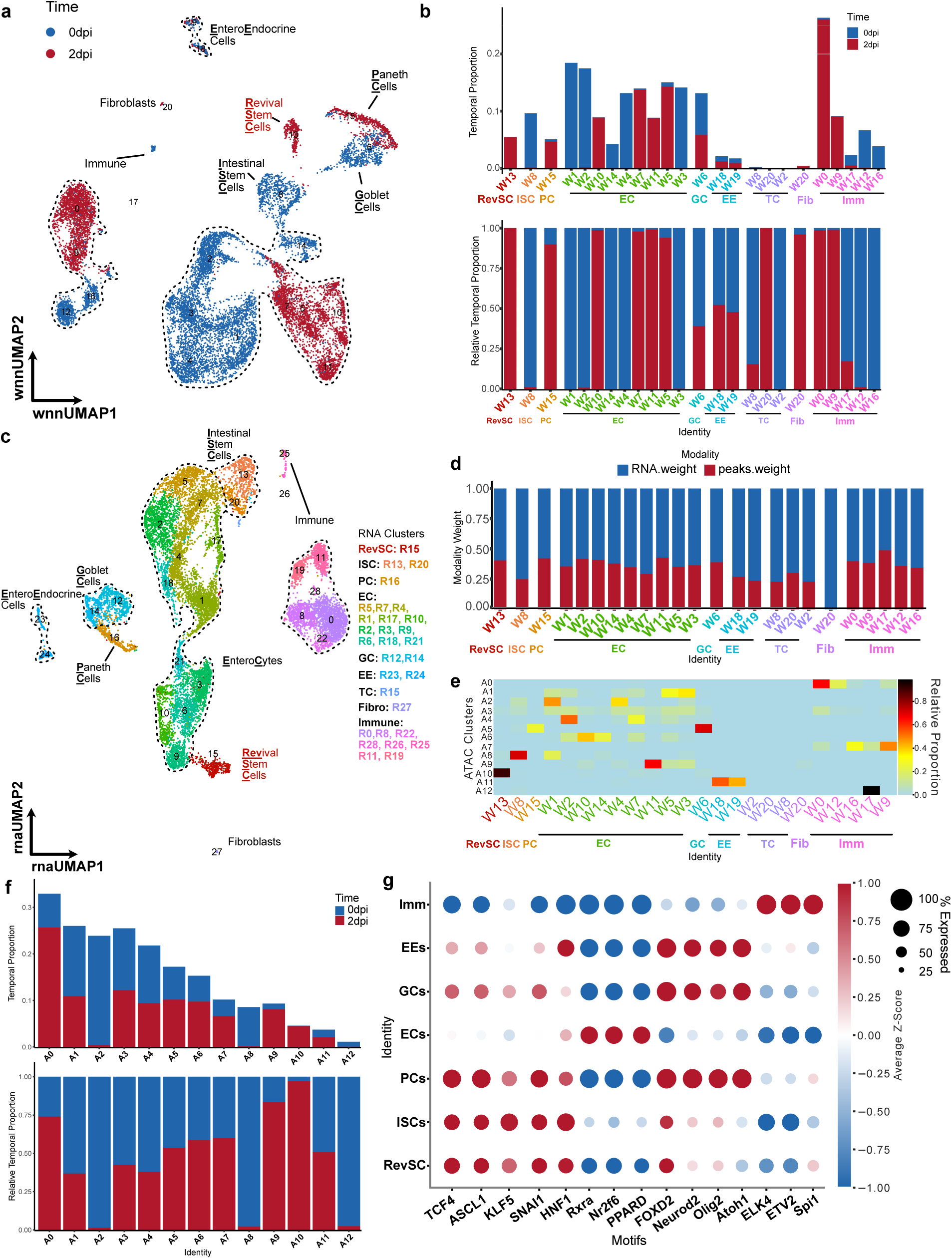
Temporal and transcriptional analysis of multiomics data set. **a)** WNN embedding of cells (1b) colored based on time point origin. **b)** Proportion of the number of cells within each identified WNN cluster relative to total number of cells from 0 dpi (blue) or 2 dpi (red) across the full embedding **(top)**. Relative proportion of number of cells within each identified WNN cluster from either 0 dpi (blue) or 2 dpi (red) relative to total number of cells from both timepoints within each cluster **(bottom)**. **c)** UMAP plot visualizing intestinal cell types generated using unsupervised clustering of single nuclei RNA-Seq data at 0 and 2 dpi. Clusters identified at higher resolutions (colour coded) were collapsed into groups corresponding to the indicated major lineages. **d)** Clusterwise median of the weighting of ATAC (red bar) and RNA (blue bar) data on the nuclei embedding for each of the indicated WNN clusters (x axis) identified in Fig 1 b. **e)** Relative proportion of nuclei (legend) from each WNN cluster (x axis) that mapped to the indicated ATAC cluster is shown as a heatmap. **f)** Proportion of the number of cells within each identified ATAC cluster relative to total number of cells from 0 dpi (blue) or 2 dpi (red) across the full embedding **(top)**. Relative proportion of number of cells within each identified ATAC cluster from either 0 dpi (blue) or 2 dpi (red) relative to total number of cells from both timepoints within each cluster **(bottom)**. **g)** Dot plot of the average ChromVAR Z-scores of each cell type for the indicated motifs. Size of the dot corresponds to the percentage of nuclei from any given cell type that that has >0 or <0 ChromVAR z-scores depending on if the average is positive or negative, respectively.

**Extended Data Fig. 2.**
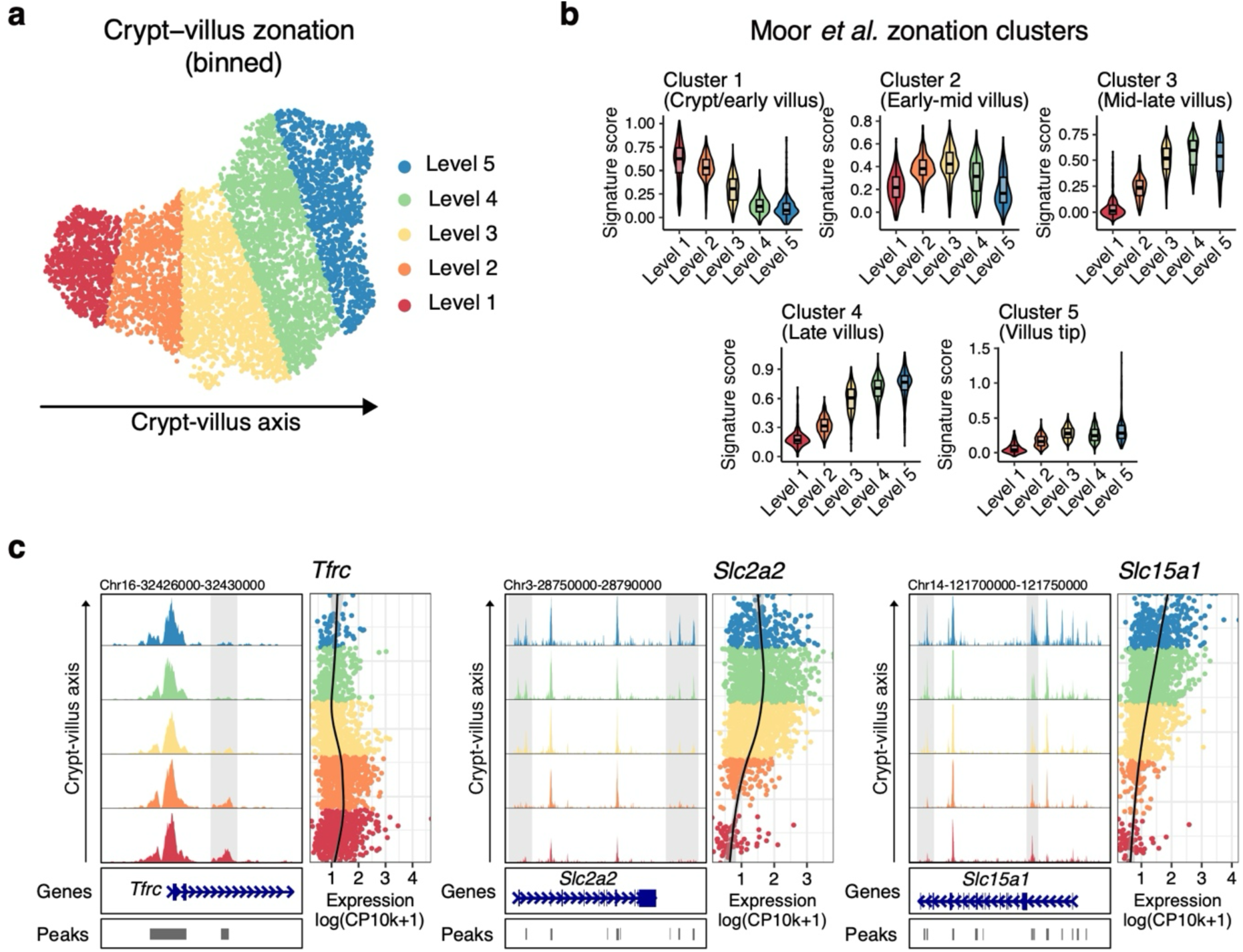
Accurate reconstruction of zonation patterns by trajectory inference. **a)** scVI-integrated UMAP embedding of 0 and 2dpi cells, discretized into five bins based on their position along the inferred trajectory. **b)** Expression levels of zonation markers defined by Moor *et al*. in each of the five discrete regions. To assess features at homeostasis, only 0 dpi cells were evaluated. **c)** Chromatin accessibility and expression evaluated at 0dpi for *Tfrc*, *Slc2a2* and *Slc15a1*. Pseudo-bulk profiles of snATAC-Seq data associated with the indicated loci are plotted as genomic traces, along with the corresponding gene structure and peak identifications (left panels). Light grey bars highlight peaks with zonation-associated accessibility. Corresponding expression is shown to the right and colour coded according to pseudotime levels. For each gene, the height of ATAC traces was set to a common range and RNA expression quantified as log-transformed UMI counts per 10k counts (log CP10k+1).

**Extended Data Fig. 3.**
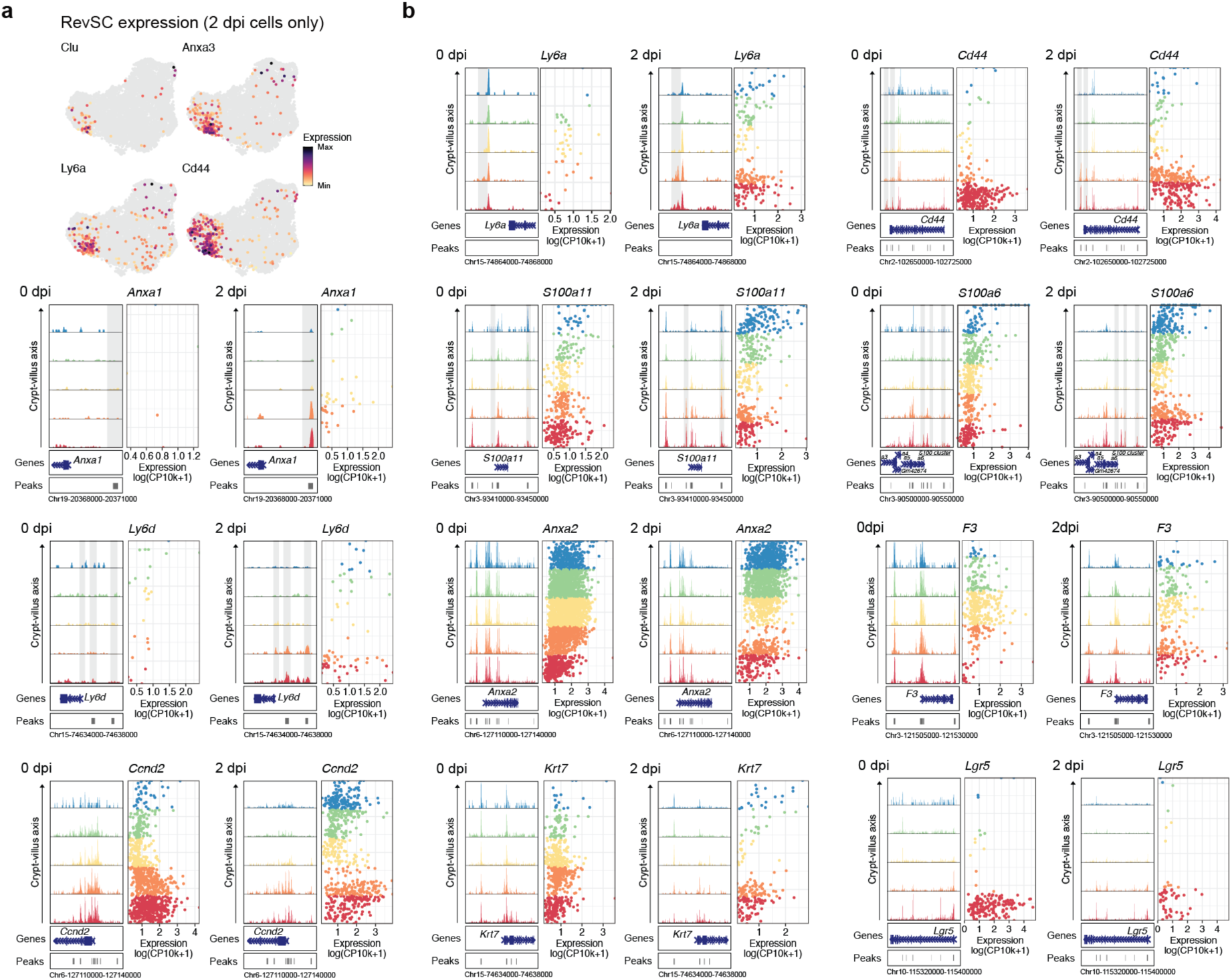
RevSC features along the crypt-villus axis. **a)** scVI-integrated UMAP embeddings showing expression of revSC associated genes. **b)** Chromatin accessibility and expression along the CV axis at 0 and 2dpi is shown for the indicated genes. Pseudo-bulk profiles of snATAC-Seq data associated with the indicated loci are plotted as genomic traces, along with the corresponding gene structure and peak identifications (left panels). Grey bars highlight injury-associated peaks. Corresponding expression is also shown, and colour coded according to levels. For each gene, the height of ATAC traces were set to a common range and RNA expression is quantified as log-transformed UMI counts per 10k counts (log CP10k+1).

**Extended Data Fig. 4.**
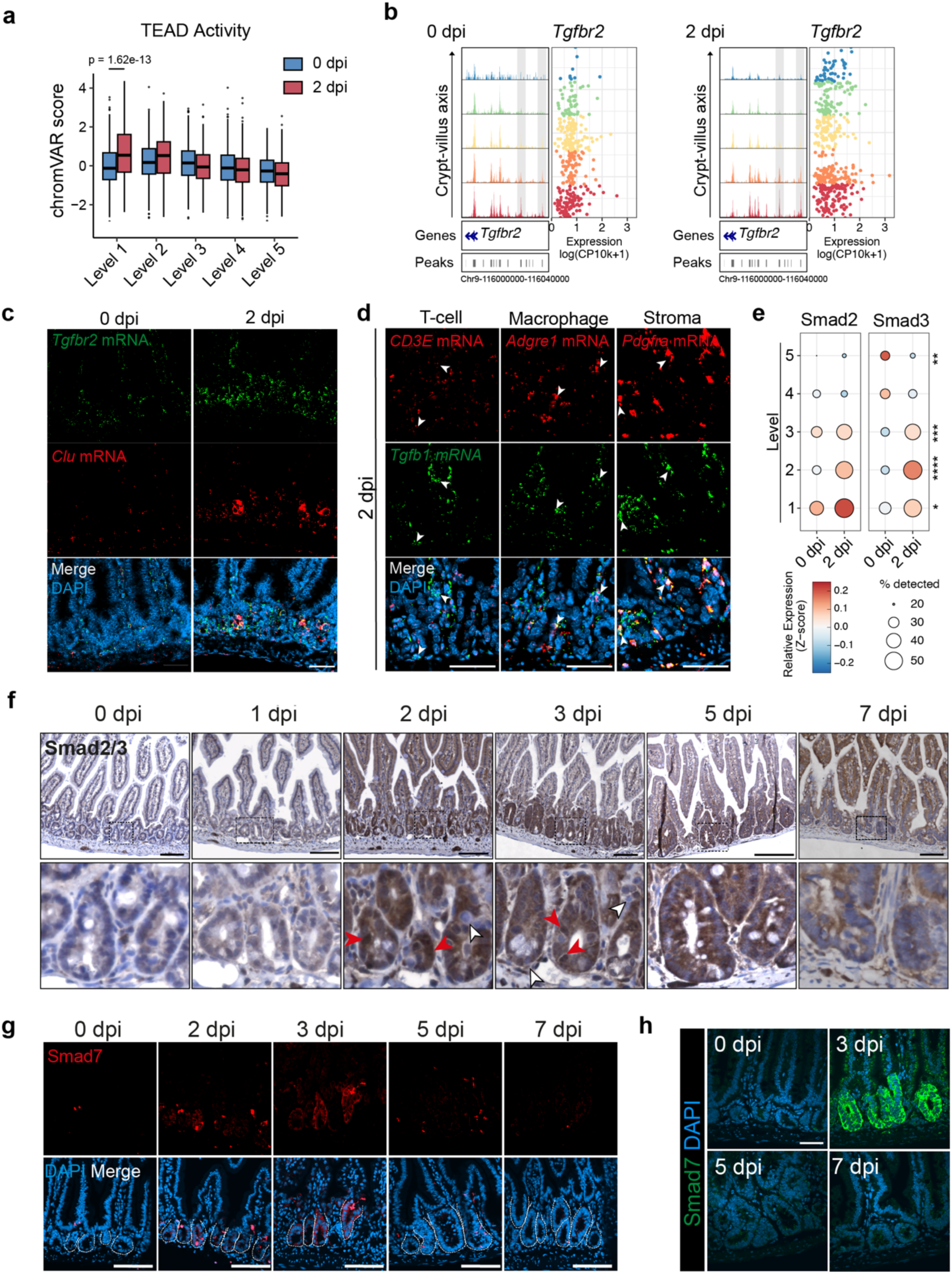
TGFβ is transiently activated in the regenerating intestine. **a)** ChromVAR activity scores for TEAD motifs in 0 and 2 dpi cells. For each cell, scores for TEAD1-4 were averaged given the similarity of their binding motif. **b)** Pseudo-bulk profiles of snATAC-Seq data associated with *Tgfbr2* in 0 and 2 dpi cells along the CV axis (left and right panels, respectively), are plotted as genomic traces, along with the corresponding gene structure and peak identifications as in Extended Data 3b. Grey bars highlight injury-associated peaks. **c)** Representative images of RNAscope for *Tgfbr2* (green) and *Clu* (red) expression in wild type mouse intestines at 0 and 2 dpi (12 Gy). Scale bar, 100 µm. n=3 mice per condition. **d)** Representative images of RNAscope of *Tgfb1* (green) expression with T-cell marker; *CD3E* (red) Macrophage marker; *Adgre1* (red) or Stromal marker*; Pdgfra* (red) at 2 dpi (12 Gy). White arrows denote co-localization. Scale bar, 50µm. n=3 mice per condition. **e)** Dot plot of median gene expression of *Smad2* and *Smad3* expression in 0 and 2 dpi nuclei along the CV axis. *p<0.05, **p<0.01, ***p<0.001, ****p<0.0001; Unpaired, two-tailed t-test. **f)** Immunohistochemistry staining of Smad2/3 protein in wild-type mice following intestinal injury at 0-, 1-, 2-, 3-, 5-, and 7-dpi (12 Gy). Boxes denote enlarged regions. Red and white arrows denote cells with or without nuclear accumulation of Smad2/3, respectively. Scale bar, 100 µm. n=3 mice per condition. **g)** Representative immunofluorescence staining of Smad7 (red) expression in wild type mice following intestinal injury at 0-, 2-, 3-, 5-, and 7 dpi (12 Gy). Dotted lines denote intestinal crypts. Scale bar, 100 µm. n=3 mice per condition. **h)** Fluorescent in situ hybridization of *Smad7* in wild type mice at 0-,3-,5-, and-7 dpi (12 Gy). Scale bar 50µm.

**Extended Data Figure 5.**
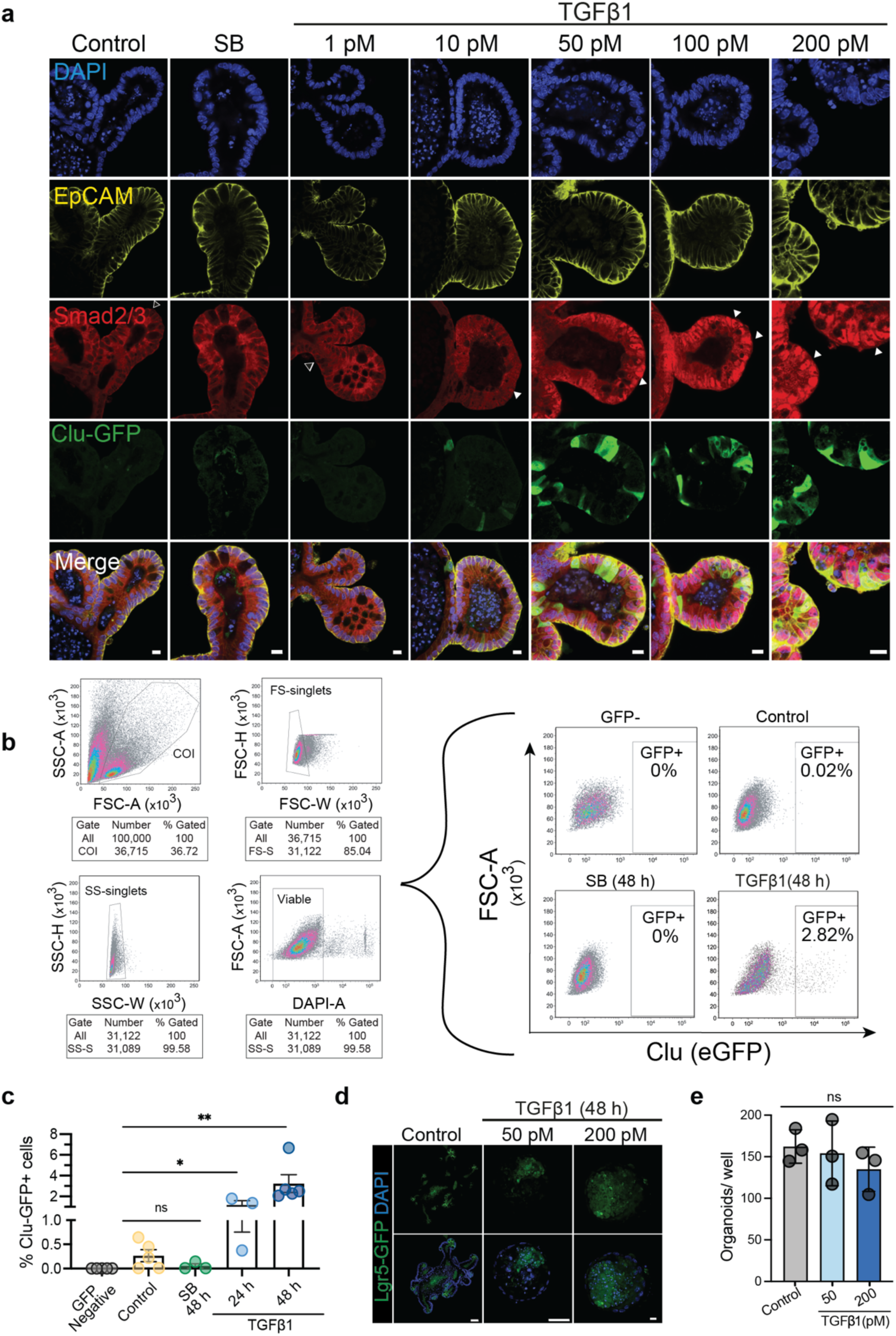
Characterizing the *in vitro* response of intestinal organoids to TGFβ pathway stimulation. **a)** Representative immunofluorescent images of Clu-GFP organoids treated with TGFβ1 or SB431542 (10 µM) for 24 hours, as indicated. Staining shows EpCam (yellow), Clu-GFP (green), and Smad2/3 (red). Nuclear Smad2/3 is indicated by solid arrows, nuclear holes indicated by open arrows. Scale bar,10µm. **b)** Gating strategy used to identify GFP+ cells isolated from organoids via flow cytometry. Plots indicate cell populations that were included based on size (singlets) and viability (DAPI-) (left). Representative plots indicating the GFP+ fraction in (1) GFP negative organoids or (2) Clu-GFP organoids that were untreated, treated with the TGFβ inhibitor, SB431542 (10 µM), or induced with TGFβ1(200 pM) (left). SS, side scatter; FS, forward scatter. **c)** Organoids expressing Clu-GFP were treated with TGFβ1 (200pM; 24 or 48 h) and the percent GFP+ cells quantified by flow cytometry. Bars represent the mean (±SEM) of n= 3 individual experiments performed in triplicate (dots). Unpaired, two-tailed t-test *p<0.05, **p<0.01. **d)** Representative IF images of Lgr5-GFP-expressing organoids treated +/− TGFβ1 (50 or 200pM; 48 h, as indicated). Scale bar, 50µm (left, middle), Scale bar, 10µm (right). **e)** Quantification of the number of organoids in culture following 24 h exposure. Organoids were treated for 24 h after 3 days of growth and the total number of organoids/well quantified. n=3 independent replicates, with 3 wells/condition. All data shown as mean ± SEM. One-way ANOVA, p= 0.5439, ns= not significant.

**Extended Data Fig. 6.**
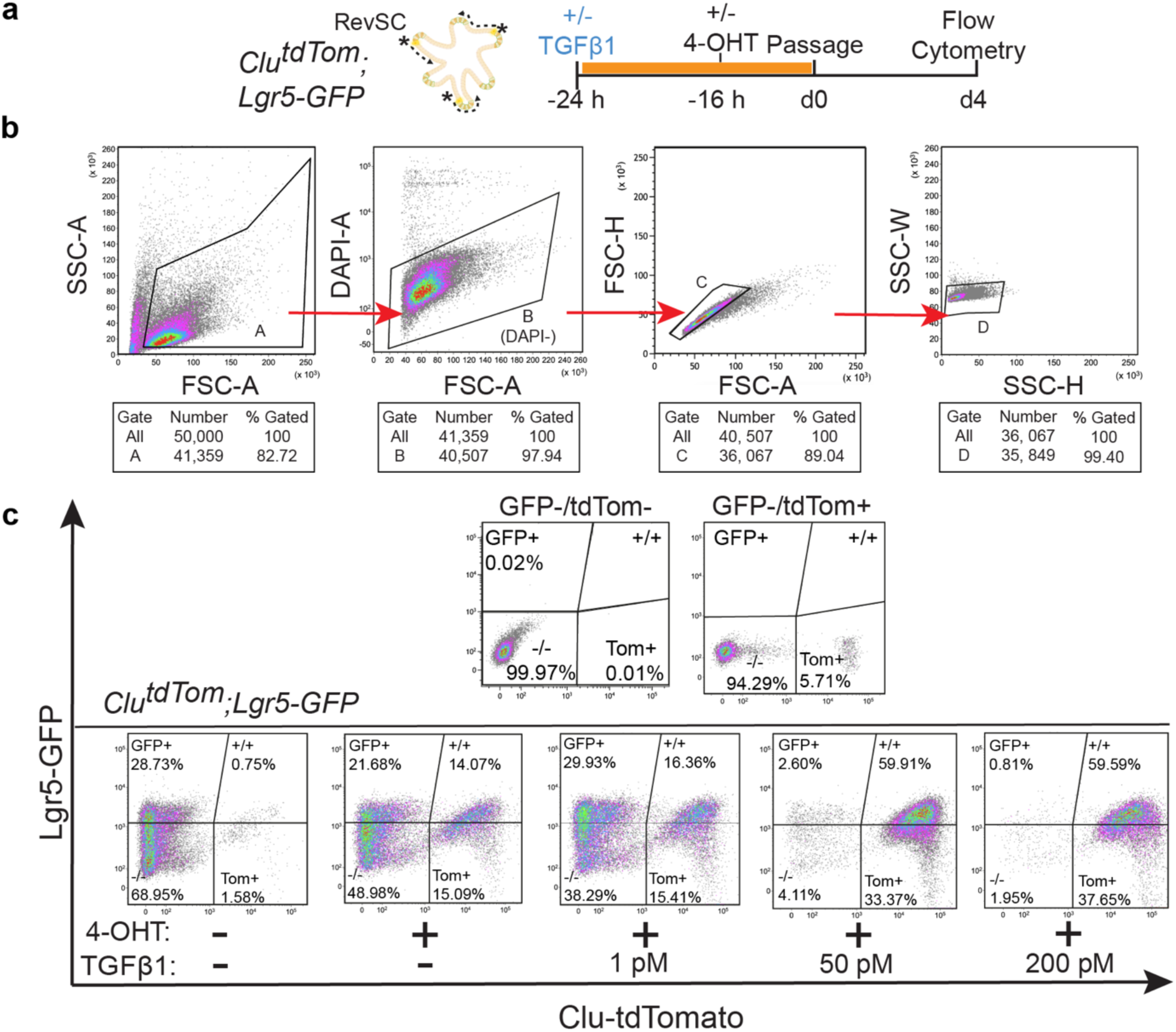
Flow cytometry analysis strategy of lineage-traced organoids. **a)** Schematic of experimental design accompanying Figure 4d. *Clu-CreERT2;lsl-tdTom;Lgr5-GFP* organoids were treated with a pulse of TGFβ (TGFβ1; 24 h) and then 4-OHT to induce recombination prior to passaging and flow cytometry. **b)** Gating strategy used to quantify cell populations isolated from organoids via flow cytometry. Plots indicate cell populations that were included based on size (singlets) and viability (DAPI-). **c)** Representative plots indicating the proportion of (1) GFP-/tdTom- (2) GFP+/tdTom-; (3) GFP+/tdTom+ and (4) GFP-/tdTom+ populations in *Clu-CreERT2;lsl-tdTom;Lgr5-GFP* organoids treated +/−4-OHT to induce lineage tracing and +/−TGFβ1 (1, 50 or 200 pM) **(bottom panels).** GFP-/tdTom- and GFP-/tdTom+ organoids were used for gating controls.

**Extended Data Fig. 7.**
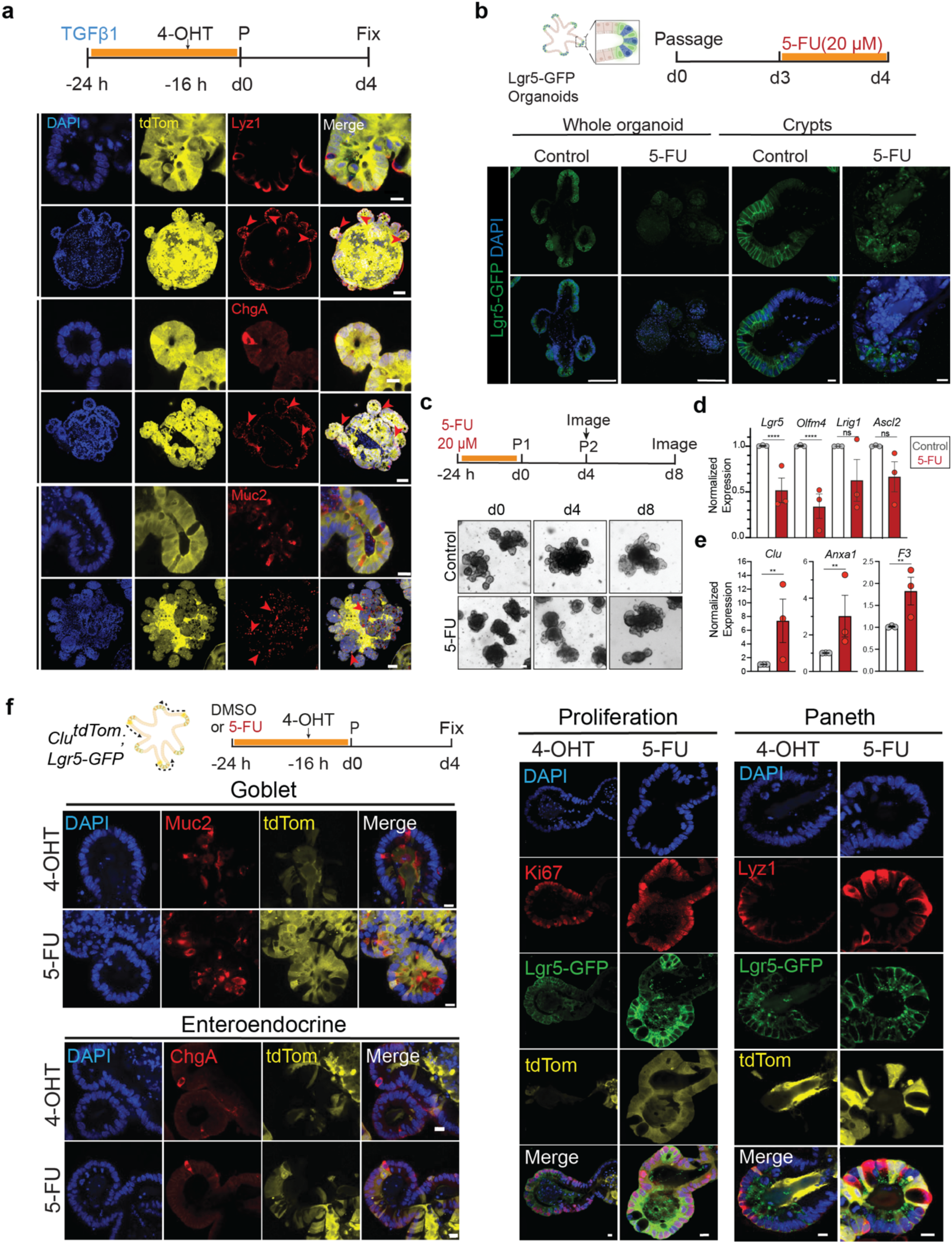
Organoids employ revSCs for functional regeneration following damage. **a)** Representative images of *Clu-tdTom;Lgr5-GFP* organoids treated with TGFβ1 (200 pM) and 4-OHT (schematic), with Paneth, Enteroendocrine and Goblet cells identified by staining for Lyz1, ChgA, and Muc2 (red), respectively. Scale bars, 10 µm (Crypt view) and 50 µm (complete organoid). Red arrows indicate cells of interest. **b)** Schematic (top) and representative IF images (bottom) of control and 5-FU-treated (20 µM; 24 h) Lgr5-GFP expressing organoids. Scale bar, 10 µm (Crypts), 50 µm (whole organoids). **c)** Schematic (top) and brightfield images (bottom) showing organoid morphology following a 24-hour pulse of 5-FU (20 µM) and after 2 successive passages. Scale bar, 10 µm. **d)** Expression of ISC signature genes in organoids treated with 5-FU (20 µM; 48 h) quantified by RT-qPCR and normalized to *Ywhaz* expression. Bars indicate mean ± S.E.M of three independent experiments performed in triplicate. Unpaired, two-tailed t-test; ***p<0.001 **e)** Expression of revSC signature genes in organoids treated with 5-FU (20 µM; 48 h), quantified by RT-qPCR, and normalized to Ywhaz expression. Bars indicate mean ± S.E.M of three independent experiments performed in triplicate. Unpaired, two-tailed t-test; ***p<0.001 **f)** *Clu-CreERT2;lsl-tdTom;Lgr5-GFP* organoids treated with 5-FU (20 µM) and 4-OHT (top) as indicated (Schematic), and lineage-traced offspring visualized after damage, as indicated. Scale bars, 10 µm.

**Extended Data Figure 8.**
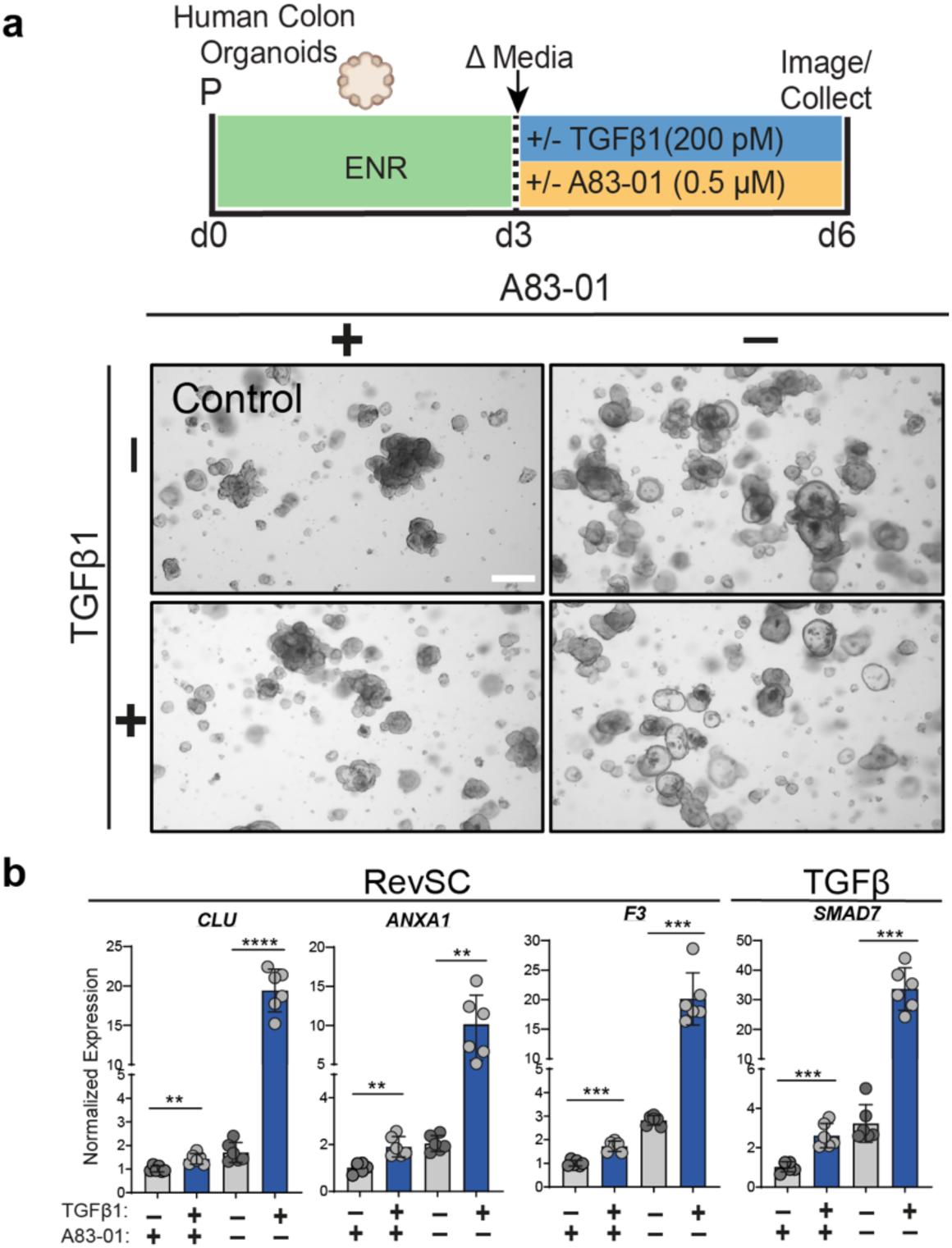
TGFβ induces a revSC response in human colon organoids. **a)** Schematic of experimental design **(top)** and representative bright-field images of human colon organoid (HCO) cultures **(bottom)**. P, passage number; d, day. +/− denotes presence or absence of TGFβ/inhibitor. Scale bar, 500 µm. **b)** qRT-PCR analysis indicating changes in revSC marker expression (*CLU, F3, ANXA1*) and *SMAD7*. Expression normalized to *GAPDH*, “+A-83-01/ −TGFβ1” represents the control. The mean (± SD) of n=6 replicates is plotted. Unpaired, two-tailed t-test *p<0.05; ** p<0.01; ***p<0.001.

**Extended Data Fig. 9.**
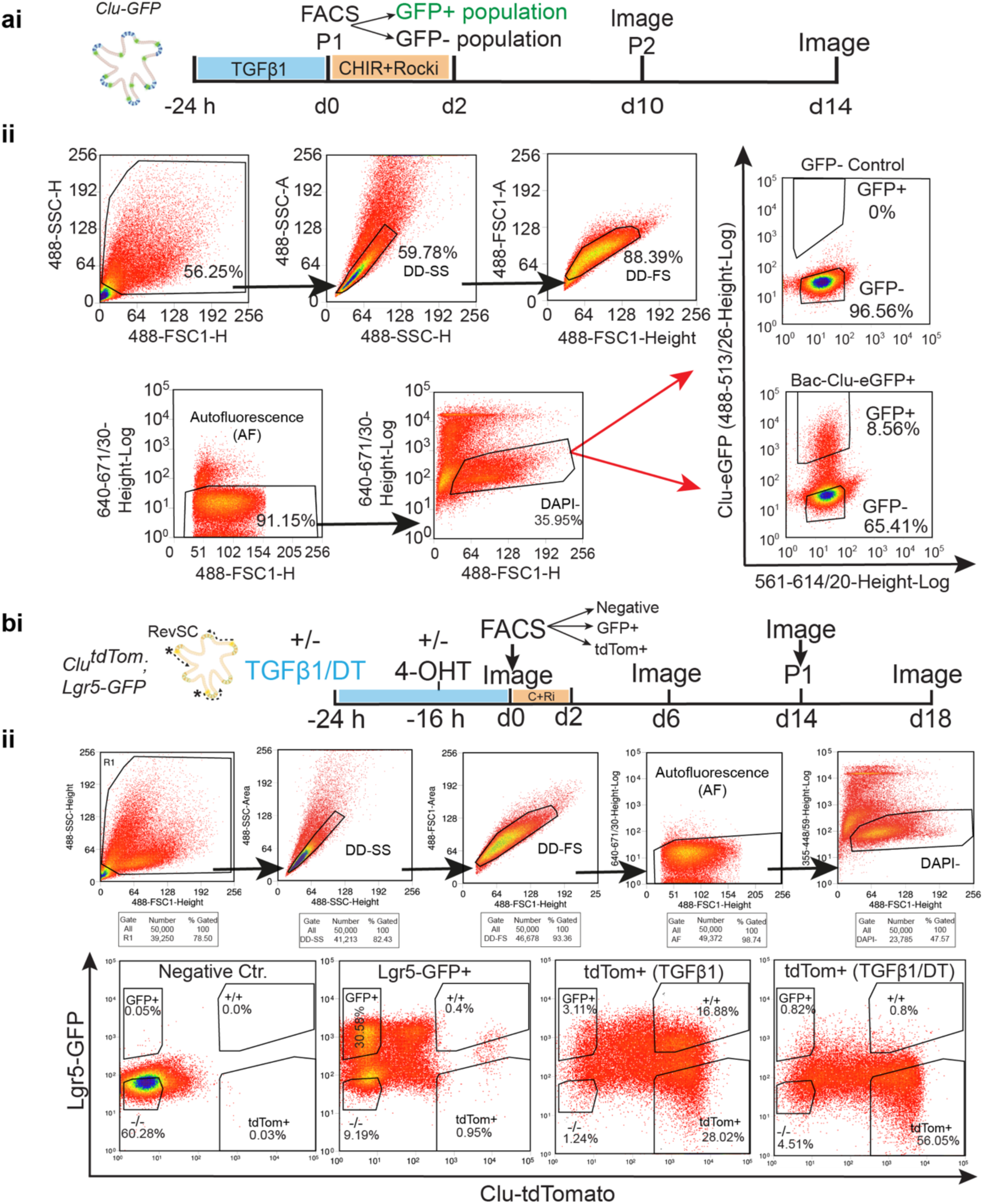
Strategy for isolating TGFβ-induced revSCs by fluorescence activated cell sorting (FACS). **a) (i)** Schematic of experimental design accompanying Figure 5a. Schematic of FACS and culture of GFP+/− cells from *Clu-GFP* organoids following treatment with TGFβ1 (24 h; 200 pM). P, passage number; d, day; h, hour; Rocki; rock inhibitor. **(ii)** Plots indicating the gating strategy used to isolate single cells from organoid cultures via FACS. Plots indicate cell populations that were included based on size (singlets) and viability (DAPI-). Representative plots of the proportion of GFP+ and GFP-cells isolated (right). Organoids lacking GFP expression were used as gating controls (top, right), and cells from *Clu-GFP* organoids were sorted into GFP+ and GFP-fractions prior to plating (right, bottom).DD, doublet discrimination; SS, side scatter; FS, forward scatter. **b) (i)** Schematic of experimental design accompanying Figure 5d. P, passage number; d, day; h, hour; Rocki; rock inhibitor; C, CHIR. **(ii)** Plots indicating the gating strategy used to isolate single cells from *Clu-CreERT2;lsl-tdTom;Lgr5-GFP* organoids via FACS. Plots indicate cell populations that were included based on size (singlets) and viability (DAPI-) (top). Representative plots showing the proportion of (1) GFP-/tdTom- (2) GFP+/tdTom-; (3) GFP+/tdTom+ and (4) GFP-/tdTom+ cells collected for plating. Organoids lacking fluorescent proteins were used as a negative control. *Clu-CreERT2;lsl-tdTom;Lgr5-GFP* organoids were treated +/− 4-OHT, +/− TGFβ1 and +/− DT as indicated in the schematic. DD, doublet discrimination; SS, side scatter; FS, forward scatter; DT, Diphtheria toxin.

**Extended Data Fig. 10.**
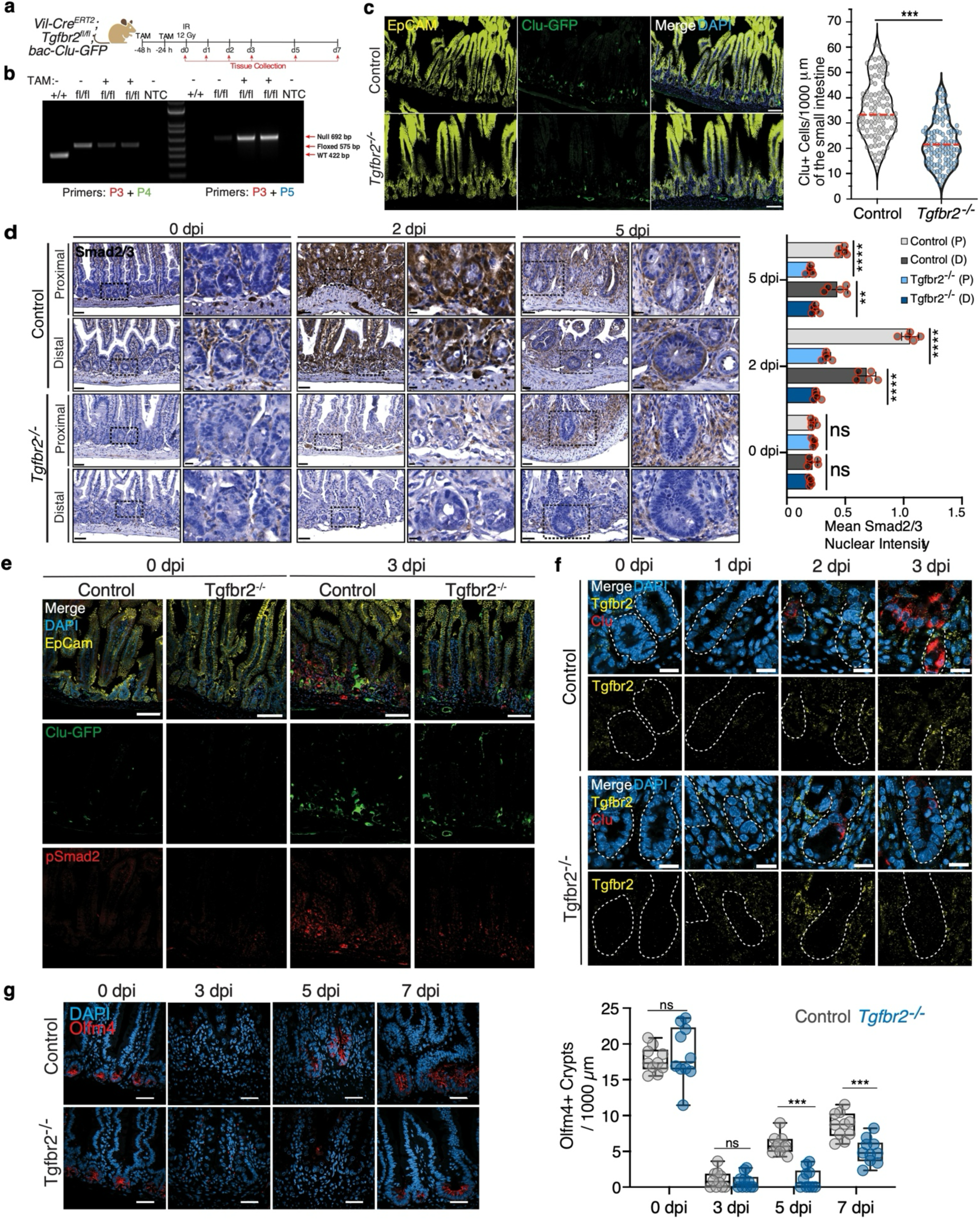
Suppressing the TGFβ pathway dampens revSC induction and delays regeneration. **a)** Schematic of *Tgfbr2* knockout in *Vil-Cre^ERT2^; Tgfbr2^fl/fl^;Clu-GFP* mice and tissue isolation following intestinal injury with 12Gy. **b)** Agarose gel electrophoresis depicting Tamoxifen-mediated Cre recombination in *Vil-Cre^ERT2^; Tgfbr2^fl/fl^;Clu-GFP* mice following tamoxifen treatment (see methods). **c)** Representative immunofluorescent images Clu-GFP (green) and EpCam (yellow) staining in *Vil-Cre^ERT2^;Tgfbr2^fl/fl^;Clu-GFP*. Control, −TAM; n=3 mice. *Tgfbr2^−/−^*, +TAM, n=3 mice (Schematic). Scale bar, 100µM **(left)**. Quantification of Clu+ cells/1000µm in the small intestine **(right)**. Clu-GFP+ cells overlapping with the epithelial-cell marker EpCAM were quantified and are plotted in a violin plot. Unpaired, two-tailed t-test; ***p<0.001, ****p<0.0001, ns=not significant. Red lines indicate median. **d)** IHC staining of Smad2/3 in Control and *Tgfbr2^−/−^* mice. Representative images depict relative expression in proximal vs. distal small intestines at 0-, 2-, and 5 dpi. Scale bar, 50 μm. Dotted boxes denote enlarged regions, scale bar, 20 μm **(right)**. Quantification of mean nuclear intensity of DAB staining per crypt **(left)**. Statistical significance using unpaired, two-tailed t-test *p<0.05, ***p<0.001, ****p<0.0001, ns=not significant. **e)** Representative immunofluorescence staining of pSmad2 (red) and Clu-GFP (green) in control or *Tgfbr2^−/−^* mice following intestinal injury at 0 and 3 dpi. *n=3* mice per condition. Scale bar, 100 μm. **f)** Representative images of RNAscope of *Clu* (red) and *Tgfbr2* (yellow) expression in Control or *Tgfbr2^−/−^* mice at 0-, 1-, 2-, and-3 dpi. *n=3* mice per condition. White dotted lines denote the crypts. Scale bar, 20 μm. g) Immunofluorescent staining denoting Olfm4 (red) expression in Control and *Tgfbr2^−/−^* mice **(left panel)**. Scale bar, 50 μm; n = 2-3 mice per condition. Quantification of Olfm4^+^ crypts per 1000 μm intestinal length **(right panel)**. Whiskers show maximum and minimum values, box extends from 25^th^ to 75^th^ percentile with median represented by the center line. Comparisons calculated by unpaired, two-tailed *t*-test ***P<0.001, ****P<0.0001, ns=not significant.

**Extended Data Fig. 11.**
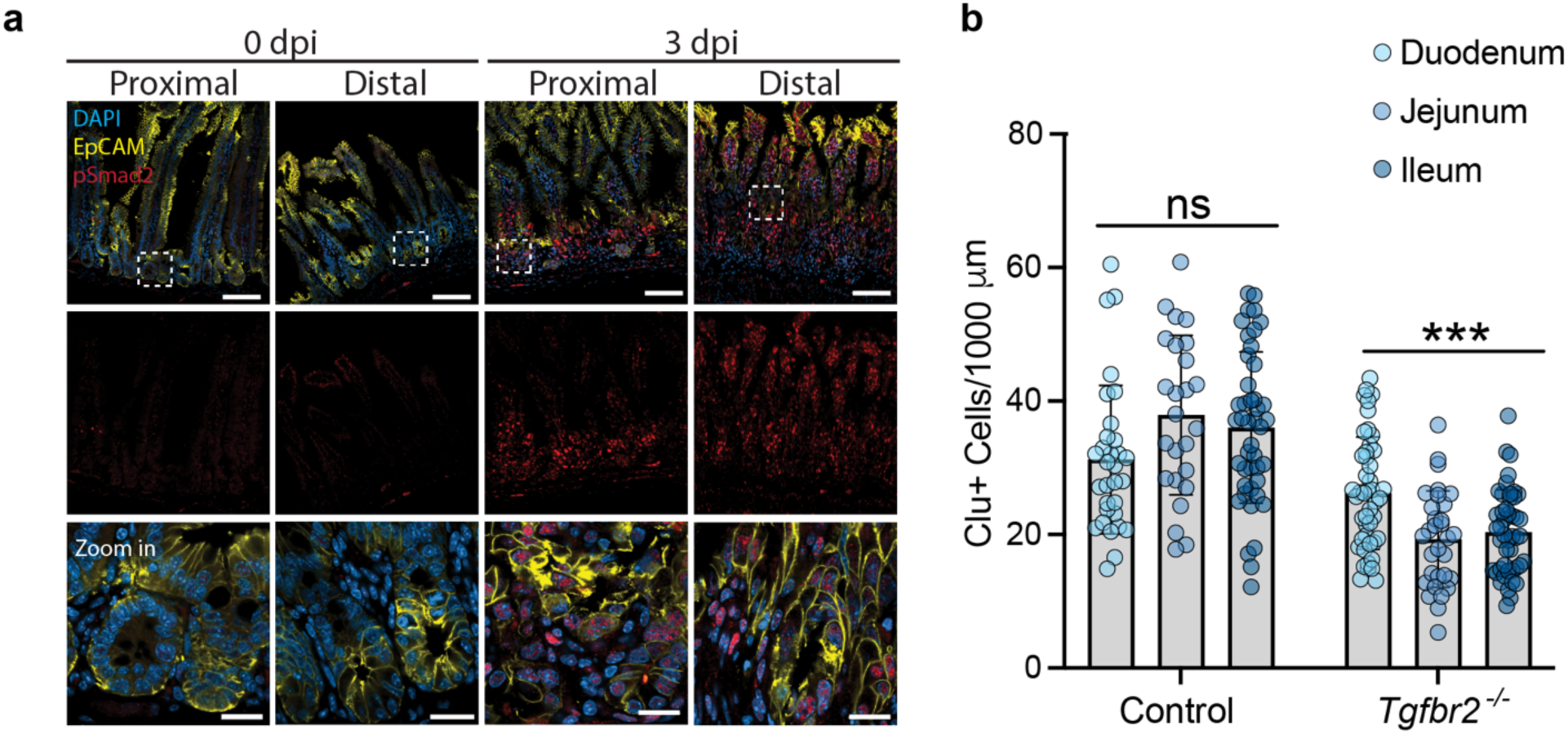
Regional Assessment of TGFβ Pathway Activation and Induction of Clu+ Cells. **a)** Immunofluorescence staining of pSmad2 in the proximal and distal small intestine of wild-type mice following intestinal injury at 0-, and 3-dpi with 12Gy. *n=3* mice per condition. Scale bar, 100 μm; scale bar in zoom in, 20 μm. **b)** Quantification of Clu+ cells/1000µm by region of the small intestine. Bars represent mean with SD. Comparison by one way ANOVA; ***p<0.001, ns=not significant.

**Extended Data Fig. 12.**
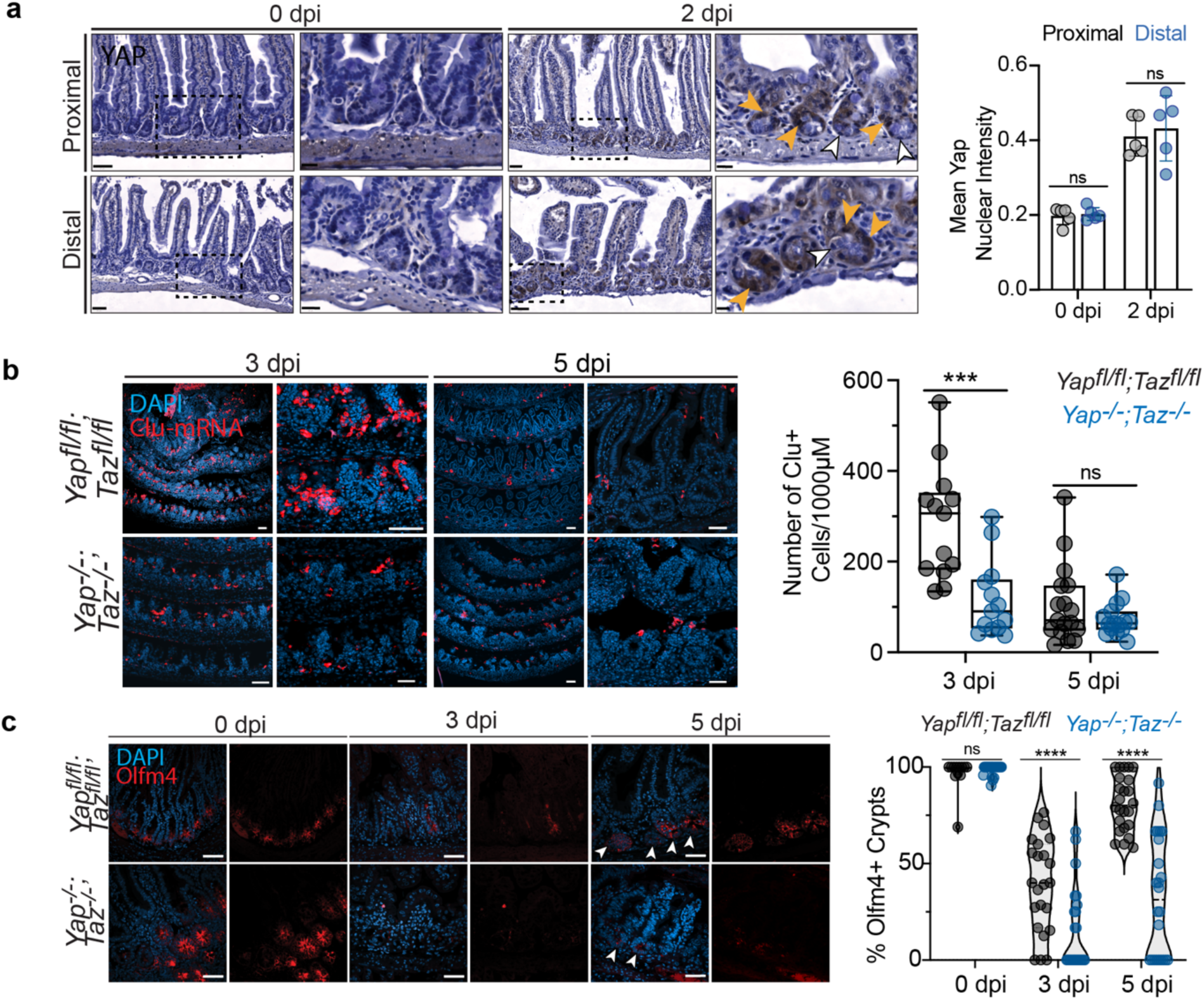
The absence of Hippo signalling reduces revSC induction and delays regeneration. **a)** Representative IHC staining of Yap in the proximal and distal regions of the small intestine at 0 and 2 dpi (12 Gy). Scale bar, 50 μm. Dotted boxes denote enlarged regions, scale bar, 20 μm. Yellow and white arrows depict Yap^+^ and Yap^−^ nuclear staining, respectively **(left)**. Quantification performed of the mean nuclear intensity of DAB staining per crypt **(right).** Statistical significance using unpaired, two-tailed t-test, ns = not significant. **b)** Representative images of RNAscope indicating *Clu* (red) expression in *Vil-CreERT2; Yap^fl/fl^*; Taz*^fl/fl^* (-TAM) and *Vil-CreERT2; Yap^−/−^*; *Taz^−/−^* (+TAM) double knockout mice at 3 or 5 dpi **(left).** Scale bar, 100 µm. *n=3* mice per condition. Quantification of Clu^+^ cells presented as box and whisker plot **(right)**. Whiskers show maximum and minimum values, box extends from the 25^th^to 75^th^ percentiles, with median represented by the center line. **c)** Immunostaining of Olfm4 expression in *Vil-CreERT2;Yap^fl/fl^*;*Taz^fl/fl^* and *Vil-CreERT2;Yap^−/−^*;*Taz^−/−^* mice at 0-, 3-, and 5 dpi (12 Gy). Scale bar, 50 μm; n = 3 mice per condition. Arrows denote intestinal crypts with re-emerging Olfm4 signal **(left)**. Quantification of Olfm4^+^ crypts per 1000 μm is presented as relative percentage **(right)**. Dotted line indicates median, Statistical analysis via unpaired, two-tailed *t*-test ***P<0.001, ****P<0.0001, ns=not significant.

